# Differentiation of human induced pluripotent stem cells into cortical neural stem cells

**DOI:** 10.1101/2022.08.19.504404

**Authors:** Alexandra Neaverson, Malin H. L. Andersson, Osama A. Arshad, Luke Foulser, Mary Goodwin-Trotman, Adam Hunter, Ben Newman, Minal Patel, Charlotte Roth, Tristan Thwaites, Helena Kilpinen, Matthew E. Hurles, Andrew Day, Sebastian S. Gerety

## Abstract

Efficient and effective methods for converting human induced pluripotent stem cells (iPSC) into differentiated derivatives are critical for performing robust, large-scale studies of development and disease modelling, and for providing a source of cells for regenerative medicine. Here, we describe a 14-day neural differentiation protocol which allows for the scalable, simultaneous differentiation of multiple iPSC lines into cortical neural stem cells (NSCs). We currently employ this protocol to differentiate and compare sets of engineered iPSC lines carrying loss of function alleles in developmental disorder associated genes, alongside isogenic wildtype controls. Using RNA sequencing (RNA-Seq), we can examine the changes in gene expression brought about by each disease gene knockout, to determine its impact on neural development and explore mechanisms of disease. The 10-day Neural Induction period uses the well established dual-SMAD inhibition approach combined with Wnt/β-Catenin inhibition to selectively induce formation of cortical NSCs. This is followed by a 4-day Neural Maintenance period facilitating NSC expansion and rosette formation, and NSC cryopreservation. We also describe methods for thawing and passaging the cryopreserved NSCs, which are useful in confirming their viability for further culture. Routine implementation of ICC Quality Control confirms the presence of PAX6-positive and/or FOXG1-positive NSCs and the absence of OCT4-positive iPSCs after differentiation. RNA-Seq, flow cytometry, immunocytochemistry (ICC) and RT-qPCR provide additional confirmation of robust presence of NSC markers in the differentiated cells. The broader utility and application of our protocol is demonstrated by the successful differentiation of wildtype iPSC lines from five additional independent donors. This paper thereby describes an efficient method for the production of large numbers of high purity cortical NSCs, which are widely applicable for downstream research into developmental mechanisms, further differentiation into postmitotic cortical neurons, or other applications such as large-scale drug screening experiments.

## Introduction

The embryonic stem cells (ESCs) found in the inner cell mass of the blastocyst-stage embryo are pluripotent, and therefore have the ability to differentiate into any of the three germ layers, producing all the cell lineages found in the adult human body. In 2007, Takahashi and Yamanaka demonstrated that somatic cells could be reprogrammed into a pluripotent state, creating induced pluripotent stem cells (iPSCs) (Takahashi et al, 2007). These cells possess many of the same properties as ESCs, importantly including their differentiation capacity and ability to self-renew, and avoid the ethical issues associated with obtaining ESCs. Since their discovery, iPSCs have been used in many areas of research including regenerative medicine, disease modelling and drug discovery. For disease modelling, their value lies in their capacity to expand continuously in culture, respond to environmental and/or genetic triggers to differentiate into cell types of interest, and the ability to derive them from the somatic cells of virtually any donor, creating patient-specific cell-based models of disease. The ease with which they can be genetically engineered also allows for the generation of isogenic models of disease carrying specific genomic alterations in an otherwise constant genetic background.

During embryonic development, ESCs differentiate, becoming increasingly lineage-committed, sequentially losing the ability to form most cell types. They pass through a multipotent stage where they are committed to differentiation into the cell types specific to one tissue or organ, for example, haematopoietic stem cells or neural stem cells (NSCs). Some multipotent stem cells persist in specialised niches in the adult body; for example, NSCs persist in the brain within the subventricular zone and the subgranular zone of the dentate gyrus in the hippocampus (Bond et al, 2016). Adult NSCs are largely quiescent, helping to avoid stem cell exhaustion, whereas embryonic NSCs are highly proliferative and expand rapidly to produce the developing CNS (Urbán and Guillemot, 2014).

In the third week of human embryonic development, the neuroectoderm forms the neural plate, which folds up and gives rise to the neural tube. NSCs line the neural tube, and will eventually give rise to all the cell types that make up the CNS. Changes in expression of developmental regulators during this period of NSC expansion or differentiation can have significant impact on brain development, and consequently on the health and survival of the child. Through genomic sequencing studies of proband cohorts, the underlying genetic causes of many rare neurodevelopmental disorders (NDDs) are being identified, and are often observed resulting from spontaneous *de novo* mutations in the genome. The Deciphering Developmental Disorders (DDD) study is a UK-wide collaborative project initiated in 2014, involving the collation of genomic and phenotypic data from children with undiagnosed developmental disorders and their parents (Wright et al, 2014). It uses large-scale genomic sequencing technologies in an attempt to identify the causative DNA variants, provide a diagnosis where possible, and identify novel genetic causes of NDDs (DDD UK, 2020). DDD-NeuGen is a large-scale project established at the Wellcome Sanger Institute in collaboration with Open Targets to capitalise on the discoveries from the DDD study, aiming to model these disorders using an NSC-based cell culture system.

The DDD-NeuGen project exploits the scalability of iPSCs to carry out large-scale differentiation of engineered disease model iPSC lines into NSCs. The loss-of-function mutations are in the same genes identified as underlying the rare developmental disorders seen in probands participating in the DDD study. The gene knockouts are created in iPSCs by inducing frame-shifting indels using CRISPR/Cas9 technology, which are then plated at low density for clonal isolation, expansion and genotyping. After colony picking and sequence based genotyping, clones carrying genotypes of interest are identified and subsequently two cell lines for each genotype are selected, expanded and differentiated into NSCs alongside unedited isogenic cell lines. The NSCs are then pelleted for downstream RNA sequencing, allowing for differential gene expression analysis, to uncover the causative mechanisms underlying each disorder and to identify similarities between disorders.

These protocols describe our methods for differentiating iPSC lines into NSCs using defined Neural Induction (NI) and Neural Maintenance stages. The 10-day NI stage uses a medium containing three different cytokines – SB431542 (hereafter referred to as SB), XAV939 (hereafter referred to as XAV) and LDN193189 (hereafter referred to as LDN), which together direct the cells towards a cortical NSC fate. SB is an inhibitor of TGFβ type 1 receptors ALK5, ALK4 and ALK7, which are involved in Activin A/Nodal signalling (Inman et al, 2002). XAV is an inhibitor of Wnt/β-Catenin-mediated transcription (Huang et al, 2009). LDN is a derivative of dorsomorphin and an inhibitor of BMP type 1 receptors ALK2 and ALK3 (Sanvitale et al, 2013). Together, inhibition of TGFβ and BMP signalling is sufficient to direct stem cells in culture down the neuroectoderm pathway (Menendez et al, 2011). Inhibition of BMP signalling combined with inhibition of Activin A/Nodal signalling is known as the dual-SMAD inhibition protocol, a well-established method for directing stem cells towards a neural precursor fate (Chambers et al, 2009). These protocols commonly use SB and Noggin, however it has been shown that LDN has a similar BMP-inhibiting effect to that of Noggin (Maroof et al 2013), and can substitute for it in these differentiations. It has also been shown that the additional inclusion of XAV in the NI medium promotes robust regionalisation of the NSCs to a dorsal cortical fate, resembling those found in the forebrain (Maroof et al, 2013). After 10 days in Neural Induction Media (NIM), the cells are passaged and re-plated in N2B27 media referred to here as Neural Maintenance Media (NMM), in which they are cultured for 4 more days before cryopreservation. The NMM encourages the formation of neural rosettes, the signature radial structures that resemble the development of the neural tube *in vitro* (Wilson & Stice, 2006). The resulting NSCs are highly enriched and will reliably express NSC markers PAX6, NESTIN and FOXG1 and lack expression of OCT4, giving highly consistent results that may vary when isogenic knockouts altering NSC development are introduced. We routinely divide each cell line into two technical replicates at the start of the NI stage, which are differentiated in parallel but kept separate throughout the protocol, thus providing independent samples; we find that these give highly concordant RNA-Seq results, demonstrating the robustness and reproducibility of the protocol.

## Materials and Equipment

### Cell Lines

All the WT parental cell lines used were generated under the Human Induced Pluripotent Stem

Cell Initiative (HipSci) (https://www.hipsci.org). All are available from ECACC with the exception of Kolf2C1_WT (https://www.phe-culturecollections.org.uk/products/celllines/generalcell/search.jsp).

1. Kolf2C1_WT, feeder-free hiPSC, male, derived from skin tissue using Cytotune 1 reprogramming on 2014-03-13. Kolf2C1_WT is a subclone of Kolf2 (HPSI0114i-kolf_2).
2. Sojd_3 (HPSI0314i-sojd_3), feeder-free hiPSC, female, derived from skin tissue using Cytotune 1 reprogramming on 2014-05-08.
3. Pelm_3 (HPSI0214i-pelm_3), feeder-free hiPSC, female, derived from skin tissue using Cytotune 1 reprogramming on 2014-05-02.
4. Podx_1 (HPSI1113i-podx_1), feeder-free hiPSC, female, derived from skin tissue using Cytotune 1 reprogramming on 2014-03-13.
5. Kucg2 (HPSI0214i-kucg_2), feeder-free hiPSC, male, derived from skin tissue using Cytotune 1 reprogramming on 2014-05-08.
6. Paim1 (HPSI0115i-paim_1), feeder-free hiPSC, male, derived from skin tissue using Cytotune 2 reprogramming.

### Cell Culture Media Recipes

#### E6 Base Media

Refer to Table 1 for concentrations and product codes of all components in E6 Base Media. Weigh out 64mg L-ascorbic acid, 1g sodium chloride and 543mg sodium bicarbonate and add to a 50ml Falcon tube. Add the mixed chemicals to 500ml of DMEM/F12. Use DMEM/F12 to wash all the residue out of the Falcon tube, ensuring all the powder is retrieved. Add the 500ml DMEM/F12 plus chemical mixture to a 1L filter unit, and add another 500ml DMEM/F12 (without chemicals). Filter through a 1L filter unit (pore size 0.2µm) to create E6 Base Media. E6 based media can be stored at −80°C for up to 12 months.

**Table 1.** Components of E6 Base Media.

| Media Component | Volume/Weight | Final Concentration |
| --- | --- | --- |
| DMEM/F12 (Life Technologies, 31330-038) | 1000ml | 100% |
| L-Ascorbic Acid (Sigma Aldrich, A8960) | 64mg | 64mg/L |
| Sodium Chloride (Sigma Aldrich, S5886) | 1g | 1g/L |
| Sodium Bicarbonate (Sigma Aldrich, S5761) | 543mg | 543mg/L |

#### Neural Induction Media

Refer to Table 2 for concentrations, volumes and product codes of all components in Neural Induction Media (NIM). Reconstitute SB431542 and XAV939 to 10mM using DMSO. Aliquots can be stored at −20°C for up to 6 months. Reconstitute LDN193189 to 1mM using DMSO. Aliquots can be stored at −20°C until the expiry date listed on the stock vial. Thaw aliquot(s) of E6 Base Media and add 50X Insulin-Transferrin-Selenium (final concentration 2X), 10mM SB43142 (final concentration 10µM), 10mM XAV939 (final concentration 2µM) and 1mM LDN193189 (final concentration 100nM). (Note: LDN193189 is light sensitive). NIM should be stored at 4°C for up to 12 days. Bring to room temperature before use. Do not warm NIM in direct sunlight or under bright lighting; if possible keep it in the dark.

**Table 2.** Components of Neural Induction Media.

| Media Component | Volume | Final Concentration |
| --- | --- | --- |
| E6 Base Media (see Table 1) | 100ml | 100% |
| Insulin-Transferrin-Selenium (100X) (Life Technologies, 41400045) | 2ml | 2X |
| SB431542 (10mM) (Abcam, ab120163) | 100μl | 10μM |
| XAV939 (10mM) (Tocris, 3748) | 20μl | 2μM |
| LDN193189 (1mM) (Cambridge Bioscience, 2092-5) | 10μl | 100nM |

#### Neural Maintenance Media

Refer to Table 3 for concentrations, volumes and product codes of all components in Neural Maintenance Media (NMM). To DMEM/F12+Glutamax (final concentration 50%), add 100X CTS N-2 (final concentration 0.5X), 50X B27 (final concentration 0.5X), 100X GlutaMAX (final concentration 0.5X), 100X MEM NEAA (final concentration 0.5X), 100X Sodium Pyruvate (final concentration 0.5X), 50mM 2-mercaptoethanol (final concentration 50µM) and 10mg/ml insulin (final concentration 2.5µg/ml). To a 500ml filter unit add a volume of Neurobasal-A medium equal to the DMEM/F12+Glutamax (final concentration 50%) and the DMEM/F12+Glutamax containing the reagents, and filter. NMM can be stored for up to 14 days at 4°C.

**Table 3.** Components of Neural Maintenance Media.

| **Media Component** | **Volume** | **Final Concentration** |
| --- | --- | --- |
| DMEM/F12+Glutamax (Thermo Fisher, 10565018) | 250ml | 50% |
| Neurobasal-A (Thermo Fisher, 12349015) | 250ml | 50% |
| CTS N-2 (100X) (Gibco, A1370701) | 2.5ml | 0.5X |
| B-27 (50X) (Gibco, 17504044) | 5ml | 0.5X |
| GlutaMAX (100X) (Thermo Scientific, 35050-061) | 2.5ml | 0.5X |
| MEM NEAA (100X) (Thermo Fisher, 11140035) | 2.5ml | 0.5X |
| Sodium Pyruvate (100X) (Sigma Aldrich, S8636) | 2.5ml | 0.5X |
| 2-mercaptoethanol (50mM) (Thermo Fisher, 31350010) | 0.5ml | 50µM |
| Insulin (10mg/ml) (Sigma Aldrich, I9278) | 125µl | 2.5µg/ml |

#### Cell Culture Materials and Labware

1. Dimethyl sulfoxide (DMSO) (Sigma Aldrich, D2650)
2. DMEM/F-12+GlutaMAX (Thermo Fisher, 10565018)
3. Dulbecco’s Phosphate Buffered Saline without MgCl_2_ and CaCl_2_ (Invitrogen – Thermo Fisher, 14190)
4. UltraPure 0.5M EDTA pH 8.0 (Invitrogen – Thermo Fisher, 15575020)
5. Essential 8 Medium (Thermo Fisher, A1517001)
6. Vitronectin (VTN-N) Recombinant Human Protein, Truncated (Thermo Fisher, A14700)
7. Recombinant Human FGF-basic (146 a.a.) (Peprotech, 100-18C)
8. ROCK inhibitor (Y-27632) (Sigma Aldrich, Y0503)
9. DNase I from bovine pancreas (Sigma, 11284932001)
10. StemPro Accutase Cell Dissociation Reagent (Thermo Fisher, A1110501)
11. 1L Filter Unit, pore size 0.2µm (Nalgene, 567-0020)
12. 500ml PES Filter Unit 0.2µM (Corning, 431097)
13. Cell Strainer 70um For 50ml Falcon Tube (Corning, 352350)
14. Corning® CoolCell® FTS30 Freezing Container for 30 × 1 mL or 2 mL Cryogenic Vials Green (Corning, 432008)
15. Cryogenic vials 1.8m (Thermo Scientific, 375418)
16. Greiner 96 well tissue culture treated black clear bottom plate (Greiner Bio-one, 655090)
17. Falcon 6 well plates (Fisher Scientific UK Limited, 10110151)

#### Immunocytochemistry Materials

1. DAPI (Biotium, 40043)
2. Donkey serum, preservative free (Bio-Rad, C06SBZ)
3. Formaldehyde, 37% (AppliChem, A0877)
4. Dulbecco’s Phosphate Buffered Saline with MgCl_2_ and CaCl_2_ (Sigma-Aldrich, D8662)
5. Triton™ X-100 (Sigma-Aldrich, 93420)

#### Primary antibodies

1. FOXG1 (EPR18987) Rabbit mAb (Abcam, ab196868)
2. OCT-3/4 (C-10) Mouse mAb (Santa Cruz Biotechnology, sc-5279)
3. PAX6 (D3A9V) XP® Rabbit mAb (Cell Signaling Technology, 60433S)

#### Secondary antibodies

1. Donkey anti-Rabbit IgG (H+L) Highly Cross-Adsorbed Secondary Antibody, Alexa Fluor 488 (Invitrogen, A-21206)
2. Donkey anti-Mouse IgG (H+L) Highly Cross-Adsorbed Secondary Antibody, Alexa Fluor 647 (Invitrogen, A-31571)

#### Flow Cytometry Materials

1. Formaldehyde, 37% (AppliChem, A0877)
2. Dulbecco’s Phosphate Buffered Saline with MgCl_2_ and CaCl_2_ (Sigma-Aldrich, D8662)
3. Fetal Bovine Serum, (Life Tech, 10270-106)
4. Triton™ X-100 (Sigma-Aldrich, 93420)
5. Primary antibodies: as above for Immunocytochemistry
6. Secondary antibodies: as above for Immunocytochemistry
7. Test tubes, polypropylene, non-sterile (Sigma-Aldrich, T1911)

#### RT-qPCR Materials

1. GenElute Mammalian Total RNA Miniprep Kit (Sigma-Aldrich, RTN70)
2. RT-PCR Grade Water (Ambion, AM9935)
3. SuperScript IV VILO Master Mix (Life Technologies, 11756500)
4. KAPA SYBR Fast Universal (Sigma-Aldrich, KK4601)
5. RT-qPCR primers (Merck Life Science UK), see Table 4

**Table 4.**
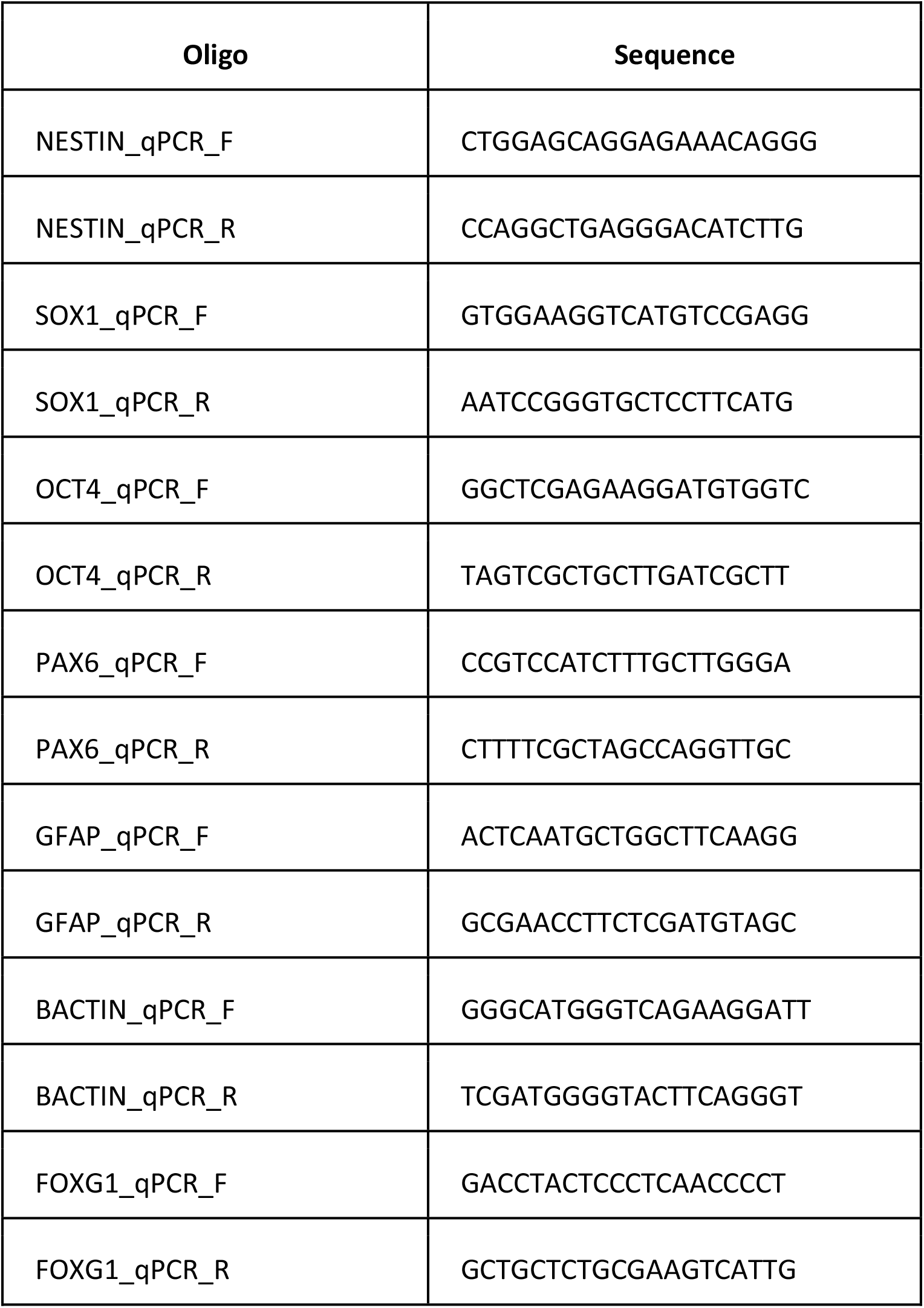
RT-qPCR Primers.

#### Equipment

1. Chemometec NucleoCounter NC-200
2. Thermo Scientific Cellomics ArrayScan XTI High Content Analysis Reader (referred to in text as Cellomics Arrayscan)
3. BD LSRFortessa Flow Cytometer

## Methods

### Generation of knockout iPSCs using CRISPR/Cas9

In order to create knockout iPSC lines, guide RNAs (gRNAs) are selected which target a conserved exon, preferably early in the protein coding sequence of a gene. For each gene target, two gRNAs are designed to account for the often unpredictable editing efficiency at different loci. The synthetic gRNA along with the Cas9 protein are delivered into the cells as a pre-complexed ribonucleoprotein via electroporation. The addition of a short single-stranded oligodeoxynucleotide of non-complementary sequence is also added to improve delivery. After a period of recovery, the cells are subcloned and up to 192 colonies picked and submitted for Next Generation Sequencing (NGS). A T7 endonuclease assay is used to assess the efficiency and inform the number of colonies needed. Clones are screened for the presence of frameshift-causing indels (insertions or deletions) and then expanded for banking. For each gene target we aim to expand both heterozygous (HT) and homozygous (HM) knockouts, as well as wild-type clones to act as controls. The wild-type clones have gone through the same treatment as the heterozygous and homozygous cells, but during the screening process are identified as non-edited. This importantly means the wildtypes should be equivalent in every other way except for the knockout mutation. After banking, the cells are sent for a second round of NGS to confirm the identity of the mutation.

The cell editing and banking work was performed by the Gene Editing facility at the Wellcome Sanger Institute.

### Pre-differentiation (up to Day −1, 2-3 weeks culture time)

The iPSC culture protocols that were followed can be found here: https://www.protocols.io/view/culture-of-established-induced-pluripotent-stem-ce-bgbwjspe

Before starting the differentiation, the knockout iPSCs are thawed and cultured on vitronectin-coated Falcon 6-well plates in E8 media (Cellular Generation and Phenotyping, 2020). Ensure that iPSCs are 70-80% confluent on Day −1. For the neural induction to work effectively, the iPSC morphology should be made up of round, dense colonies of cells with minimal spontaneous differentiation present in the culture. It may be necessary to passage the iPSCs several times to achieve this morphology.

As part of the DDD-NeuGen pipeline, two wild-type (WT1 and WT2), two heterozygous (HT1 and HT2) and two homozygous (HM1 and HM2) isogenic knockout iPSC lines are differentiated simultaneously alongside the parental iPSC line (Kolf2C1_WT). Each cell line is split into two technical replicates during the Neural Induction passage on Day −1. Together these cell lines constitute one ‘gene set’, allowing for large-scale, simultaneous culture of cells with different knockout genotypes for the same disease gene. We aim to cryopreserve 5 NSC vials (each containing 2.5×10^6^ cells) and create 3 cell pellets for each technical replicate; in order to achieve these numbers, we differentiate 4 wells (of a Falcon 6-well plate, 9.6 cm^2^ per well) per replicate. As a starting point, we recommend expanding each iPSC line to 12 wells across two Falcon 6-well plates ready for neural induction. For simplicity, the protocol below describes the steps stating the volumes required per well of a Falcon 6-well plate. Please note that all centrifugation steps were done at room temperature unless otherwise indicated.

### Neural Induction (Day −1 to Day 10)

This section describes how to differentiate an iPSC line into NSCs over a period of 10 days. During the Neural Induction passage, each cell line is divided up into technical replicates which are then kept separate for the rest of the differentiation.

1. Coat the required number of Falcon 6-well plates with vitronectin diluted 1:50 in DPBS(-/-) and incubate for at least 2 hours at 37ºC.
2. Aspirate media from all the wells of one iPSC line and wash with 2ml DPBS(-/-) per well. Aspirate DPBS(-/-) and add 1ml accutase to each well. Incubate for 3-6 minutes at 37ºC, checking after 3 minutes to see if cells have begun to detach.
3. Add 4ml E8 media per well, pipetting up and down using a stripette to dislodge the cells. It is important to break down the iPSC colonies into a single cell suspension. Collect all cells into a 50ml Falcon tube.
4. Determine the cell count per ml using a Chemometec NucleoCounter NC-200, or other appropriate cell counter. Calculate the volume of cell suspension required to seed 2×10^6^ live cells per well of a 6-well plate (approximately 200,000 cells per cm^2^). We recommend seeding 8 wells per cell line (4 per technical replicate).
5. Isolate the required volume into a 15ml Falcon tube. If you are planning RNAseq or qPCR analysis of the cell line, isolate the required volume of the suspension into separate 15ml Falcon tube(s); 2 million cells per cell pellet is recommended. Centrifuge all cells for 5 minutes at 200 rcf.
6. Make up seeding media by supplementing E8 media with ROCK inhibitor (Y-27632) to a final concentration of 10μM. ROCK inhibitor promotes survival of cells after single cell dissociation. Seeding media can be made up in bulk to have enough for the whole gene set. Make up enough seeding media to give 2ml per well to be re-plated into.
7. Aspirate supernatant and re-suspend the cells in 2ml E8 plus ROCK inhibitor per 2×10^6^ cells.
8. Quickly remove vitronectin from the newly coated plates and transfer 2ml cell suspension into each well. Gently shake back and forth and side to side, ensuring that the cells become evenly distributed.
9. Transfer cells to a 37°C, 5% CO_2_ tissue culture incubator overnight, gently agitating the plate once more.
10. To create cell pellets, aspirate supernatant media and re-suspend the cell pellet with DPBS(-/-) and divide between Eppendorf tubes. Microcentrifuge the tubes for 30 seconds to pellet the cells, then aspirate supernatant and snap-freeze. Store snap-frozen cell pellets at −80 °C.
11. Repeat Steps 2-10 for each additional cell line.
12. 24 hours after plating, the cells should have attached in a uniform sheet across the well. From this point, the cells are prone to peeling. When aspirating, do not touch the cell layer at any point. Media should be dispensed gently on the side of the well, never directly onto the cells.
13. Aspirate the media from the wells and gently wash with 2ml DPBS(-/-) per well.
14. Aspirate the DPBS (-/-) and slowly dispense 2ml NIM per well, down the side of the well. Return the cells to the incubator. This is Day 0 of Neural Induction.
16. Media change the cells every 24 hours with 2-3ml NIM per well, until Day 10.
17. The cells should form a neuroepithelial sheet around Day 3 (see Figure 2 for examples of morphology). Towards the end of the differentiation period, proliferation is high, the media may be very yellow, and there is an increased risk of the neuroepithelial sheet peeling off on later days, so take care to dispense media very slowly. If there is excessive peeling, it might be necessary to passage the cells ahead of Day 10.

**Figure 1.**
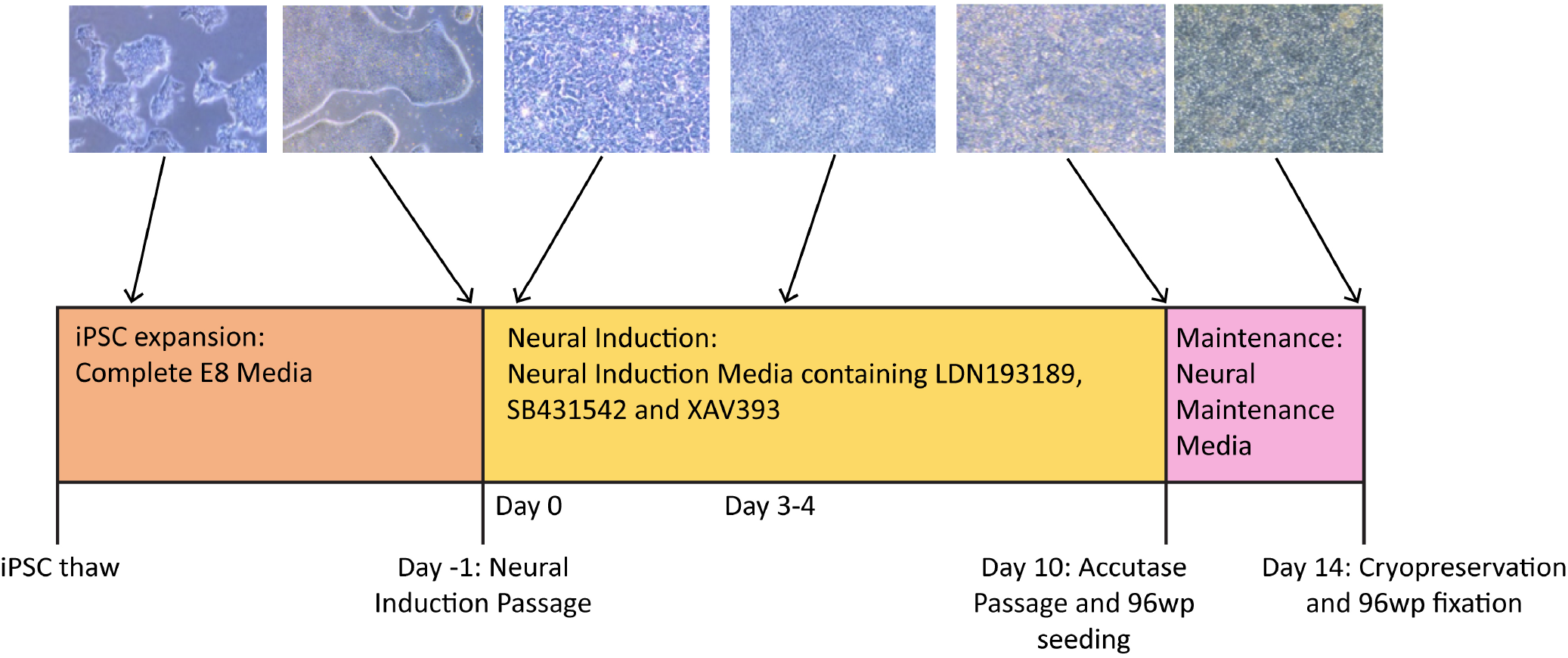
Timeline of the neural differentiation protocol. iPSCs are thawed in Complete E8 media, then expanded to the desired number of wells before the Neural Induction passage (Day − 1). On Day 0, the medium is switched to NIM (Neural Induction Media). Media change with NIM takes place daily until Day 10, when the cells are passaged with accutase and simultaneously plated to 96-well plates for immunocytochemistry. By Day 10, the cells should have neural stem cell identity. During this passage, the cells are re-plated in NMM (Neural Maintenance Media) on both plate types. The cells grow in NMM until Day 14, with media change taking place every other day. On Day 14, the cells are cryopreserved in NMM + 10% DMSO, and those growing on the 96-well plates are fixed in preparation for antibody staining.

**Figure 2.**
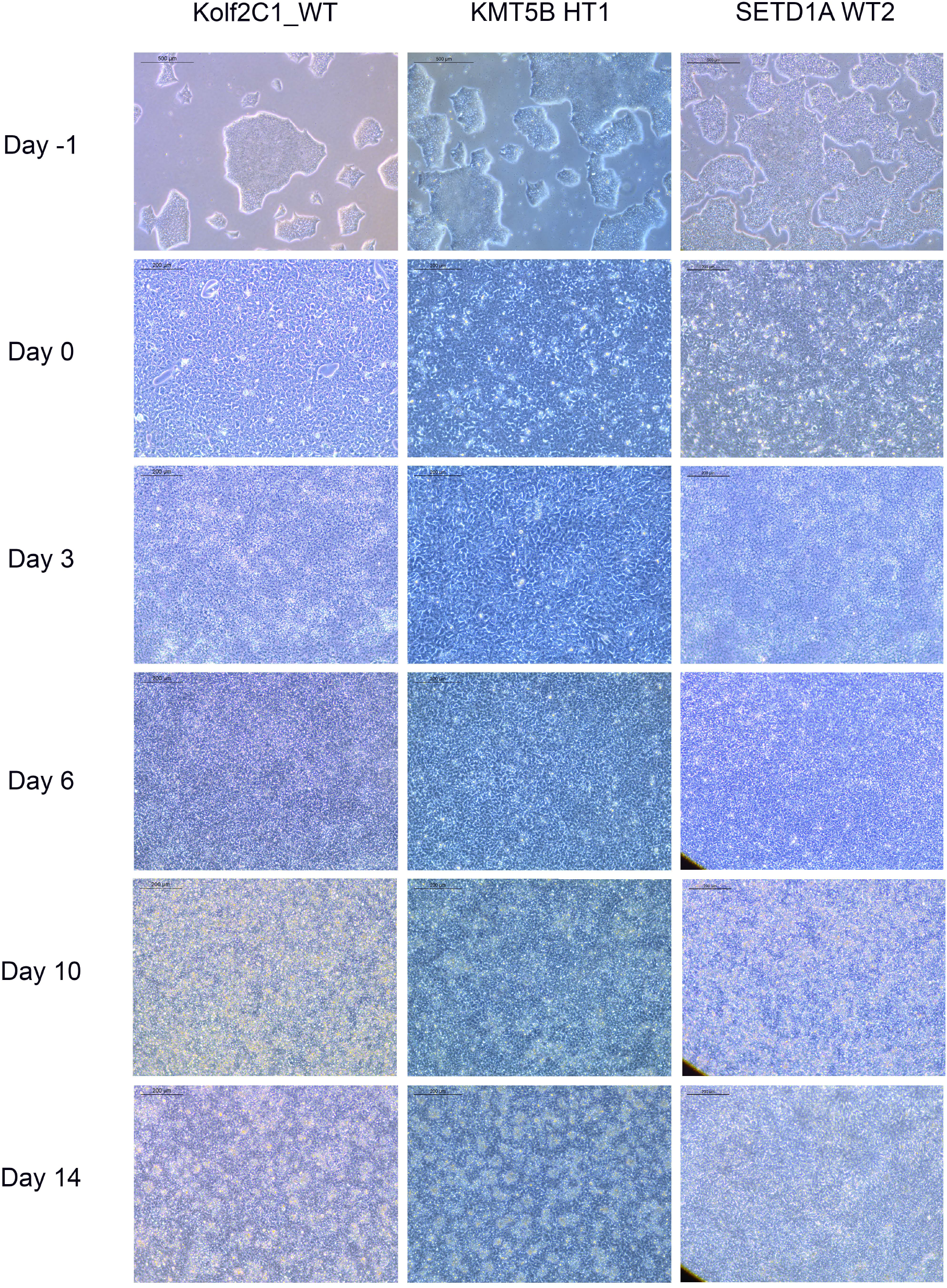
Representative cell lines at each stage of the 14 day neural differentiation protocol. Displayed are images of Kolf2C1_WT, a heterozygous KMT5B knockout (KMT5B HT1) and a wild-type line from the SETD1A gene set (SETD1A WT2). Note the differences in morphology between each of the cell lines, but all of these examples constitute acceptable morphology at each stage. For example, SETD1A WT2 has clear neural rosettes on Day 14, whereas the others do not. Scale bars: Day −1, 500μm (5x). All other images: 200μm (10x).

### Accutase Passage of Neuroepithelial Sheet (Day 10)

18. Coat Falcon 6-well plates with 1:100 vitronectin in DPBS (-/-) and incubate for at least 2 hours at 37ºC. Coat the same number of wells and plates as for Neural Induction. It is recommended that the cells are also plated onto 96-well plates for the ICC QC assay during this passage; if so, coat 96-well clear-bottomed tissue culture plate(s) with 1:100 vitronectin in DPBS(-/-). Note 1: For the ICC QC Assay to be effective, a negative control in the form of undifferentiated iPSCs is required. It is recommended that you have a minimum of 1 well of 70-80% confluent, good quality iPSCs growing in preparation for plating onto the 96-well plate. Note 2: Because we use rabbit primary antibodies for both PAX6 and FOXG1, these markers can not be used in the same sample, so our protocol requires two independent plates for each gene set. We recommended coating one column per differentiated cell line, per plate, plus the parental iPSC line as a negative control. The number of plates to coat will depend on the number of cell lines and antibody species you are using.
19. Supplement NMM with ROCK inhibitor (Y-27632) to a final concentration of 10μM. Prepare 4ml NMM with ROCK inhibitor per well, plus an additional 3ml per cell line for re-plating onto the 96-well plate(s) for ICC.
20. Working with one cell line at a time, aspirate NIM from cells and gently wash with 2ml DPBS (-/-) per well. Aspirate DPBS and add 1ml accutase to each well. Incubate for 3-6 minutes at 37ºC, checking after 3 minutes to see if the cells are detaching.
21. While the cells are incubating, prepare the wash buffer containing DMEM/F12+Glutamax and DNAse I at a final concentration of 30μg/ml. Prepare a minimum of 4ml wash buffer per well. This should be prepared fresh for each cell line to ensure optimal DNAse activity.
22. After incubation, add 4ml wash buffer to each well, pipetting up and down using a stripette to dislodge the cells. Collect all wells for one technical replicate into a 50ml Falcon tube. Mix the cell suspension thoroughly and aliquot into 15ml Falcon tubes of equal volume for centrifugation. It is recommended that one technical replicate per cell line is selected for plating onto 96-well plate(s) for ICC. For the selected technical replicate, isolate 2.5ml of the cell suspension into a separate 15ml Falcon tube, and centrifuge all tubes together for 5 minutes at 200 rcf.
23. Aspirate the supernatant from the cells for re-plating onto the 6-well plates, and resuspend them in 4ml NMM per well to be re-plated into. Since this passage is 1:1, this should be the same number of wells as in the Neural Induction stage.
24. Quickly remove the vitronectin from the newly coated plates and transfer 4ml cell suspension into each well. Agitate the plate back and forth and side to side, to ensure the cells become evenly distributed.
25. Transfer plates to a 37°C 5% CO_2_ tissue culture incubator overnight, gently agitating the plate once more.
26. For the cells isolated for ICC, aspirate the supernatant and resuspend in 3ml NMM plus ROCK inhibitor. This will allow for enough cells at the correct density to seed to 12 wells of a 96-well plate. Aspirate the vitronectin from one column on the coated 96-well clear-bottomed tissue culture plate(s). Seed 200μl each to 6 wells of one column on (each) 96-well plate. Transfer the plate(s) to a 37°C 5% CO2 tissue culture incubator overnight, gently agitating the plate(s) once more.
27. Repeat steps 20-26 with the remaining cell lines.
28. To plate iPSCs to the 96-well plates for ICC, detach iPSCs using EDTA diluted 1:1000 in DPBS(-/-), harvest and gently resuspend in 10ml E8 media. Isolate 500μl of the cell suspension into a 15ml Falcon tube, and dilute using an extra 1ml of E8 media. This will allow for enough cells at the correct density to seed to 12 wells of a 96-well plate. Aspirate the vitronectin from one column on the coated 96-well clear-bottomed tissue culture plate(s). Seed 100μl each to 6 wells of one column on (each) 96-well plate(s). Transfer the plate(s) to a 37°C 5% CO2 tissue culture incubator overnight, gently agitating the plate(s) once more.

### Neural Stem Cell Maintenance Culture (Day 11 – Day 13)

29. Culture the cells for three days, changing the media on Day 11 and Day 13 with 4ml NMM (Neural Maintenance Media) per well. During this period, the cells should proliferate and reach a morphology of small round/oval cells. Neural rosette structure should be visible when viewing the cells under a microscope with a 10x or 20x objective.
30. Culture the cells on the 96-well plates for three days, media changing the NSCs with 200μl NMM on Day 11 and Day 13, and media changing the iPS cells 100μl E8 daily. The cells should be fixed using formaldehyde on Day 14 (see section 3.9).

### Cryopreservation of Neural Stem Cells (Day 14)

31. Prepare neural freeze media by combining NMM (90%) and DMSO (10%).
32. Aspirate the NMM from the wells of your plates, working with one cell line at a time. Wash gently with 2ml DPBS(-/-) per well. Aspirate DPBS and add 1ml accutase per well and incubate in a humidified incubator at 37°C for approximately 4 minutes, checking after 2 minutes to see if the cells are detaching.
33. Add 4ml DMEM/F12+Glutamax to each well, pipetting up and down with a stripette to break up the cell layer.
34. Pass the cells through a 70μm filter, collecting all the cells from each technical replicate into separate 50ml Falcon tubes.
35. Determine the cell count using an appropriate cell counter. Use a suitable method to determine cell viability, for example trypan blue staining. NSCs should be frozen at 2.5×10^6^ live cells per vial. Isolate the required amount of cells for the number of vials you wish to freeze, dividing the volume evenly between 15ml Falcon tubes. Note: If you are planning RNA-Seq or qPCR analysis of the cell line, isolate the required volume of cell suspension into separate 15ml Falcon tube(s) for making pellets. A minimum of 2×10^6^ cells per cell pellet is recommended.
36. Centrifuge the cells for 5 minutes at 200 rcf.
37. Remove the supernatant from the cells to be cryopreserved. Resuspend in 1ml neural freezing media per 2.5×10^6^ cells, then divide the cell suspension between 1.8ml cryovials, adding 1ml to each cryovial. Place the cryovials into a CoolCell or other appropriate freezing container, then transfer to a −80°C freezer. Cells should be transferred to liquid nitrogen storage after 24 hours.
38. To create cell pellets, aspirate the supernatant and re-suspend the cell pellet in DPBS(-/-). Divide the cell suspension between 2ml Eppendorf tubes. Microcentrifuge the tubes for 30 seconds to pellet the cells, then aspirate the supernatant and snap-freeze on dry ice. Store snap-frozen cell pellets at −80°C.

### Thawing Neural Stem Cells

39. Coat Falcon 6-well plates with 1:100 vitronectin in DPBS(-/-) and incubate for at least 2 hours at 37ºC. NSCs are frozen at a density which allows for plating 1 vial to 1 well of a 6-well plate.
40. Prepare NMM supplemented with ROCK inhibitor (Y-27632, to a final concentration of 10μM) and bFGF (final concentration 20ng/ml). Prepare 4ml per vial to be thawed.
41. Partially thaw cells in a 37ºC water bath. Add 1ml DMEM/F12+Glutamax to the cryovial dropwise and transfer the partially thawed cells to a fresh 15ml Falcon tube containing a further 7ml DMEM/F12+Glutamax. Use another 1ml DMEM/F12+Glutamax to wash the cryovial to ensure all cells are collected.
42. Centrifuge the cells for 5 minutes at 200 rcf.
43. Aspirate the supernatant and gently resuspend the cell pellet in 4ml NMM plus ROCK inhibitor and bFGF. Note: If cell numbers in the frozen vial were unknown, it is recommended to perform a cell count before plating and aim to seed at a density of 0.15-0.25×10^6^ cells/cm^2^ (1.5-2.5×10^6^ cells per well of a 6-well plate).
44. Remove vitronectin from the prepared plate and replace with 4ml cell suspension at the correct density.
45. Gently agitate the plate back and forth and side to side, to ensure cells become evenly distributed.
46. Transfer plates to a 37°C 5% CO_2_ tissue culture incubator overnight, gently agitating the plate once more.
47. 24 hours after plating, media change with 4ml fresh NMM supplemented with 20ng/ml bFGF.
48. Media change every other day with NMM supplemented with 20ng/ml bFGF. The cells should be confluent enough to passage after 4-6 days of culture.

### Passaging Neural Stem Cells

49. Coat Falcon 6 well plates with 1:100 vitronectin in DPBS(-/-) and incubate for at least 2 hours at 37ºC. Since this passage is 1:1, the same number of wells/plates will be required as was used for thawing NSCs.
50. Aliquot NMM into a Falcon tube and supplement with ROCK inhibitor (final concentration 10μM) and bFGF (final concentration 20ng/ml). Prepare 4ml per well to be re-plated into.
51. Aspirate the media from the cells, and wash with 2ml DPBS(-/-).
52. Aspirate DPBS and add 1ml accutase per well and incubate in a humidified incubator at 37°C for approximately 4 minutes, checking after 2 minutes to see if the cells are detaching.
53. Add 4ml DMEM/F12+Glutamax per well, gently pipette up and down to dislodge cells and collect into a 50ml Falcon tube. Note: If cells remain very clumpy despite accutase treatment, it is possible to filter them using a 70μm filter into a 50ml Falcon tube.
54. Determine the cell count using an appropriate cell counter. Calculate the volume of cell suspension required to seed 2×10^6^ cells per well of a 6-well plate (0.2×10^6^ cell per cm^2^). If you have additional cell suspension remaining, it may be possible to expand the culture at this stage.
55. Centrifuge the cells for 5 minutes at 200 rcf.
56. Aspirate supernatant and gently re-suspend the pellet in 4ml NMM supplemented with ROCK inhibitor and bFGF per well to be seeded.
57. Remove vitronectin from the prepared plate and replace with 4ml of the cell suspension.
58. Gently agitate the plate back and forth and side to side, to ensure cells become evenly distributed.
59. Transfer plates to a 37°C 5% CO_2_ tissue culture incubator overnight, gently agitating the plate once more.
60. 24 hours after plating, media change with 4ml fresh NMM plus 20ng/ml bFGF.

### Immunocytochemistry Quality Control Assay

A detailed version of our ICC Protocol can be found here: https://www.protocols.io/view/differentiation-of-human-induced-pluripotent-stem-bgbxjspn

The ICC assay is used to determine the success of neural differentiation by estimating the frequency of expression of NSC markers. The stem cell marker OCT4 is used to identify the fraction of non-differentiated iPS cells. On day 14 of neural differentiation, cells are fixed using 4% formaldehyde at room temperature for 20 minutes, followed by three washes of Dulbecco’s Phosphate Buffered Saline with MgCl_2_ and CaCl_2_ (DPBS+/+). (Pause point: Fixed cells can be kept in DPBS+/+ at 4°C for up to 10 days before proceeding to the next step). Cells are then incubated in 10% donkey serum diluted in DPBS+/+ with 0.1% Triton-X100 for 2 hours as a combined permeabilisation and blocking step. Primary antibodies are diluted as above, but in 1% donkey serum to the desired dilution (1:200 for PAX6, 1:2000 for FOXG1, 1:100 for OCT4), and cells are incubated with the diluted antibodies for 1 hour at room temperature. At the end of the incubation time, cells are washed twice with DPBS+/+. Secondary antibodies (Alexa Fluor 488 and Alexa Fluor 647) and DAPI are diluted 1:1000 in 1% donkey serum. Cells are incubated with secondary antibodies for 1 hour at room temperature, in the dark. At the end of the incubation time, cells are washed three times with DPBS+/+ and then analysed using a Cellomics Arrayscan. The Arrayscan uses an automated protocol to identify DAPI-stained cell nuclei and measure the percentage of the DAPI-positive cell population which is also exhibit positive staining for the assay markers (PAX6, FOXG1, OCT4) which are all nuclear markers. Noise and background staining are normalised using a no-primary antibody staining control.

### Flow Cytometry

Flow cytometry was used to quantify marker-positive cell fractions (PAX6, OCT4, FOXG1) and compare to the same data acquired through ICC. Two vials each of Kolf2C1_WT, SETD1B WT and SETD1B HT knockout NSCs were thawed onto 1:100 VTN, grown for 4 days in NMM supplemented with 20ng/ml bFGF. Kolf2C1_WT NSCs were passaged using EDTA and grown in E8 media for 4 days until 70% confluent. Both cell types were then dissociated using accutase, and fixed in suspension using 1% formaldehyde in DPBS +/+. 1×10^6^ cells per cell line per experimental condition were fixed, leaving the cells to stand for 30 minutes at room temperature in suspension with 1% formaldehyde. The cells were washed twice by centrifuging for 3 minutes at 300rcf, aspirating supernatant, then resuspending in 2ml FACS buffer (5% FBS in DPBS +/+).

The cells were permeablised using 2ml DPBS+/+ with 0.1% Triton-X for 30 minutes, before centrifuging for 3 minutes at 300rcf, then washing with 1ml FACS buffer as described. The cells were then divided up into their respective experimental conditions (PAX6, OCT4, FOXG1 and no-primary control), then left suspended in FACS buffer for 30 minutes to block. The cells were then centrifuged for 3 minutes at 300 rcf, the supernatant was aspirated, then resuspended in 100µl primary antibody solution. Antibody concentrations were as follows: PAX6 1:400, OCT4 1:400, FOXG1 1:4000 in FACS buffer (see section 3.9 for details of antibodies used). The cells were incubated with the primary antibodies for 1 hour at room temperature, before centrifugation for 3 minutes at 300 rcf. The cells were then resuspended in 100µl secondary antibody solution, made up of FACS buffer plus AF467 and AF488 both at 1:1000 (see section 3.9 for details of antibodies used). They were then incubated for 1 hour at room temperature in the dark. Cells were centrifuged for 3 minutes at 300 rcf, then washed with 1ml FACS buffer, centrifuged again and resuspended in 350µl FACS buffer and transferred into FACS tubes, before analysis on the BD LSRFortessa flow cytometer using FACS Diva software. Cells were vortexed immediately before analysis, and run through the analyser using the fastest speed setting. FACS Diva was used to categorize cell populations based on their expression of each marker.

### RT-qPCR

For carrying out gene expression analysis of the MED13L gene set, RT-qPCR was performed on the gene set (containing 2 HT, 2 HM and 2 WT lines) after differentiation into NSCs. Gene expression levels were calculated as fold change in expression compared to undifferentiated Kolf2C1_WT iPSCs. Cell pellets were collected at the end of neural differentiation for MED13L, at the cryopreservation stage. Kolf2C1_WT iPSC pellets were created from 70% confluent wells. RNA was extracted using a GenElute Mammalian Total RNA Miniprep Kit, as per manufacturer’s instructions. Total RNA was quantified using a NanoDrop One (Thermo Fisher), and 400ng was added to a reverse transcription reaction with 4µl Superscript IV VILO Master Mix and made up to 16µl with RT-PCR grade water. Reverse transcription was performed using a thermal cycler (MJ Research PTC-240) to produce cDNA. The resultant cDNA was diluted in 815ul of Milli-Q water, then amplified and quantified through qPCR with SYBRGreen on the Stratagene Mx300p qPCR machine. The housekeeping gene ACTB(β-actin) was used as a control. Milli-Q water was used in the place of cDNA as a control for contamination. Each sample was tested in triplicate. Primer sequences are provided in Table 4. Gene expression fold change was quantified using the 2^-ΔΔ*Ct*^ method, first normalising against the Ct values for β-actin, then calculating relative fold change compared to Kolf2C1_WT iPSCs.

RT-qPCR was also performed on the HipSci lines Sojd_3, Podx_1, and Pelm_3 in addition to the Kolf2_C1 (which has been used throughout the experiments). Cell pellets for each line were collected prior to neural differentiation (on Day −1) and at the end of the neural differentiation (Day 14), as described above. The same procedure as above was followed, with the exception that fold-change in expression was calculated relative to the iPSC sample for each cell line.

### RNA-Seq methods

RNA extraction from frozen cell pellets containing up to 2×10^6^ cells was performed using manufacturer’s protocols for their RNeasy QIAcube kit on a QIAcube automated purification system (Qiagen), and RNA sequencing libraries were prepared using established protocols. Briefly, library construction (poly(A) pulldown, fragmentation, 1st and 2nd strand synthesis, end prep and ligation) was performed using the ‘NEB Ultra II RNA custom kit’ (New England Biolabs) on an Agilent Bravo WS automation system. We performed indexed, multiplexed sequencing on the Novaseq 6000 system (S4 flow cell, Xp workflow; Illumina), collecting approximately 30M X 100 base, paired-end reads. The sequencing data were de-plexed into separate CRAM files for each library in a lane. Adapters that had been hard-clipped prior to alignment were reinserted as soft-clipped post alignment, and duplicate fragments were marked in the CRAM files. The data pre-processing, including sequence QC, and STAR alignments were made with a custom Nextflow pipeline, which is publicly available at https://github.com/wtsi-hgi/nextflow-pipelines/blob/rna_seq_5607/pipelines/rna_seq.nf, including the specific aligner parameters. We assessed the sequence data quality using FastQC v0.11.8. Reads were aligned to the GRCh38 human reference genome (Ensembl GTF annotation v91). We used STAR version STAR_2.6.1d (Dobin *et al*., 2013) with the --twopassMode Basic parameter. The STAR index was built against GRCh38 Ensembl GTF v91 using the option -sjdbOverhang 75. We then used featureCounts version 1.6.4 (Liao, Smyth and Shi, 2014) to obtain a count matrix.

Genes with no count or only a single count across all samples were filtered out. The counts were normalised using DESeq2’s median of ratios method (Love, Huber, & Anders, 2014). A pseudocount of one was added to the count matrix followed by log2 transformation. Plots for marker expression and replicate scatterplots were created with R (R Core Team, 2019) using ggplot2 (Wickham, 2016), ggpubr (Kassambara, 2020) and ggbeeswarm (Clarke & Sherrill-Mix, 2017) packages.

## Results

### Neural Differentiation Results

Our neural differentiation protocol provides a 14-day method for the generation of human neural stem cells from iPSCs, with two distinct phases, neural induction and neural maintenance. Using this approach, we have been able to differentiate multiple iPSC gene sets (containing heterozygous knockout, homozygous knockout and wild-type cell lines) successfully into NSCs. We typically expect to achieve a cell count of 2-4×10^7^ cells from 4 wells of a 6-well plate, on Day 14. The morphology observed throughout neural induction is consistent; Figure 2 shows examples of typical cell morphology at each stage of differentiation, including the slight variations that still constitute acceptable morphology.

On Day −1, iPSCs should be of good quality, displaying rounded colonies with smooth edges and spontaneous differentiation should be minimal or absent (Figure 2, “Day −1”). They should be approximately 70-80% confluent to ensure that cell count is optimal to begin neural induction. The day after carrying out the neural induction passage, the cells will have formed a monolayer (Figure 2, “Day 0”), and the E8 medium is replaced by NIM. Between Day 1 and Day 4, the cells will become smaller and take on a ‘frogspawn-like’ morphology, where individual, round cells and their nuclei are visible at 10x magnification (Figure 2, “Day 3”). From Day 5-10, they will continue to shrink and proliferate, until individual cells are no longer visible at 10x magnification (Figure 2, “Day 6”). On Day 10, you should have a relatively homogeneous layer of NSCs (Figure 2, “Day 10”). These are then passaged with accutase and re-plated in NMM. During Days 10-14, the NSCs mature and form the characteristic polarised structures known as neural rosettes (Figure 2, “Day 14”). These may be visible under a light microscope at 10x magnification as in Figure 2, SETD1A WT2, or they may be harder to spot; after staining with NSC-specific antibodies, rosettes are much more easily observed as a rose or star-shaped pattern (for example, as in Figure 8A).

### Troubleshooting Neural Differentiation

#### Potential Problem 1: iPSC Quality

iPSCs are poor quality and/or not 70-80% confluent on the intended day of neural induction passaging. They may be overgrown (see Figure 3A) or not forming distinct colonies, appearing spiky and uncompact (Figure 3B). As a rule, 70-80% confluence should be the aim in order to optimize growth rate after seeding. The iPSCs should be passaged until they improve in quality, before starting differentiation. Passaging with a higher split ratio may help to improve iPSC compaction. Additionally, try to limit the number of times the iPSCs are pipetted, during the EDTA passage, in order to preserve the clumps of cells, and minimise the number of single cells generated.

**Figure 3.**
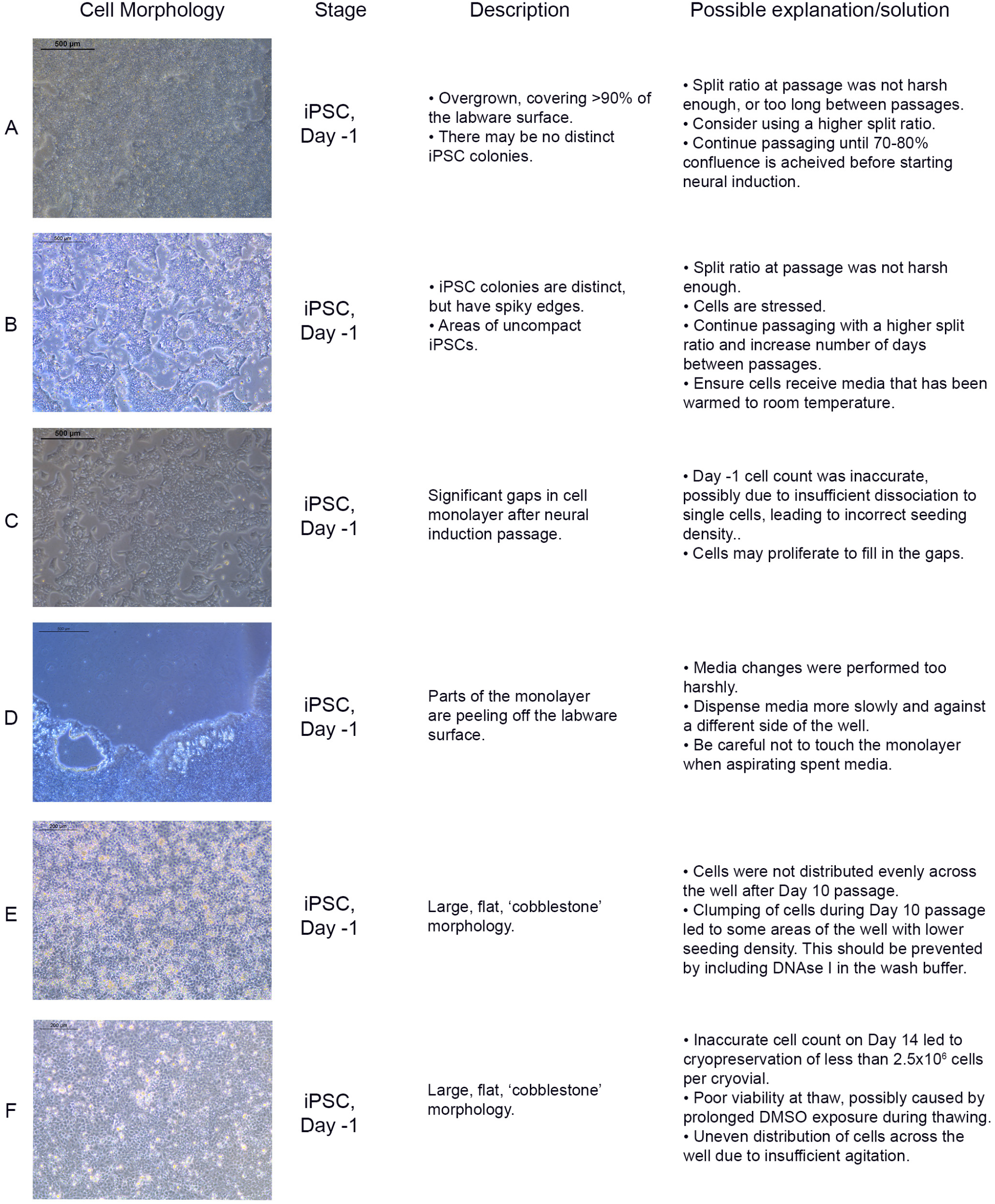
Common differentiation problems. Examples of common issues at different stages of neural differentiation, their morphological appearance, possible explanations as to why they may occur, and potential corrective actions.

#### Potential Problem 2: Low density on Day 0

On Day 0 of neural induction, the cells are not forming an even monolayer, but instead have gaps between them (see Figure 3C). This may be due to an uneven distribution of cells, from insufficient agitation of the plate after passaging, or the number of live cells seeded may be less than 2×10^6^, which could be due to an inaccurate cell count before seeding. If the gaps are minor (i.e. approximately 90-95% confluent), the cells should proliferate enough to fill the gaps within 1-2 days. Larger gaps will result in poor NSC differentiation and increased appearance of undesired neural derivatives such as neural crest cells.

#### Potential Problem 3: Peeling of monolayer

During neural differentiation, the cells become very fragile and are prone to detaching from the labware surface, particularly from the edges of the wells (see Figure 3D). If they start to peel off, reduce the speed of the media change, and dispense medium onto the opposite side of the well, ensuring that the medium runs down the side of the well rather than directly onto the cells. If the cells are becoming increasingly fragile, consider passaging on Day 9 instead of Day 10.

#### Potential Problem 4: Morphology issues

After the Day 10 passage, NSCs appear large and exhibit a flat, square morphology (see Figure 3E). The NSCs may have been distributed unevenly across the well, particularly if this morphology is only seen in part of the well. This morphology is commonly seen if the cells become clumpy during the passage; inclusion of DNAse in the wash buffer should prevent this. In our experience, cells can recover from this between Day 11-14, and even if morphology remains the same, the cells usually still express NSC markers.

#### Potential Problem 5: Lack of dissociation after accutase treatment

NSCs are not dissociating when incubated with accutase for >5 minutes. When this issue occurs, it is usually seen during the Day 10 passage. If this occurs, dispensing the wash buffer quickly onto the cells multiple times may encourage them to detach. If this does not work, gently scrape off as many cells as possible from the labware surface using a stripette. Inclusion of DNAse I in the wash buffer should prevent excessive clumping from the release of DNA from lysed cells. Consider increasing the accutase incubation time (but ensure it is kept under 10 minutes), however be aware that this may increase cell death.

### Immunocytochemistry Results

On Day 10, the NSCs are passaged using accutase and seeded both to 6-well plates, and to 96-well plates for ICC. These grow alongside each other and receive the same media (NMM). The 96-well plates are fixed on Day 14, in parallel with the cryopreservation of those cultured on 6-well plates, and should therefore be representative of the banked cells. The ICC is an important QC step of the pipeline, checking the percentage of the cell population expressing PAX6 and FOXG1 (we use >80% positively expressing as a passing cut-off) and the absence of expression of the iPSC marker, OCT4. Images are collected and expression data is quantified using the Cellomics Arrayscan, however similar results can be obtained using any fluorescence microscope. The main advantages of the Cellomics Arrayscan approach are the ability to collect a large number of images relatively quickly with little user input, and the ability to generate quantitative gene expression data through unbiased counting, achieved by automation. Figure 8A shows the ICC results of the MED13L gene set, where in this case the mutation has not visibly impacted differentiation and all NSCs display high levels of PAX6 staining, including the heterozygous knockouts. The use of ICC also allows for the visual inspection of the effectiveness of the neural differentiation, and a judgement to be made on the quality of the resulting cells. Figure 4 shows PAX6 expression data for 16 gene sets, demonstrating the consistency between gene sets successfully differentiated over a 1-year period. Of note is the observation that lines performing poorly by this metric are more likely to be knockout samples, rather than wild-type or unedited parental lines, and so may represent a biological effect of the gene knockout. Of all cell lines differentiated in this time period, the mean percentage of PAX6-positive cells was 85.6% (data not shown). Figure 5 shows some examples of suboptimal outcomes observed during ICC, which can then be flagged for closer inspection and possible exclusion. This includes circular clumps of PAX6-negative cells, patchy PAX6 expression, and residual OCT4-positive iPS cells. These outcomes may indicate a failure of differentiation, rather than a technical issue with the ICC process, and may be due to a biological (potentially knockout-related) effect. If this is thought to be the true outcome of the differentiation, we may undertake a repeat differentiation to confirm the finding.

**Figure 4.**
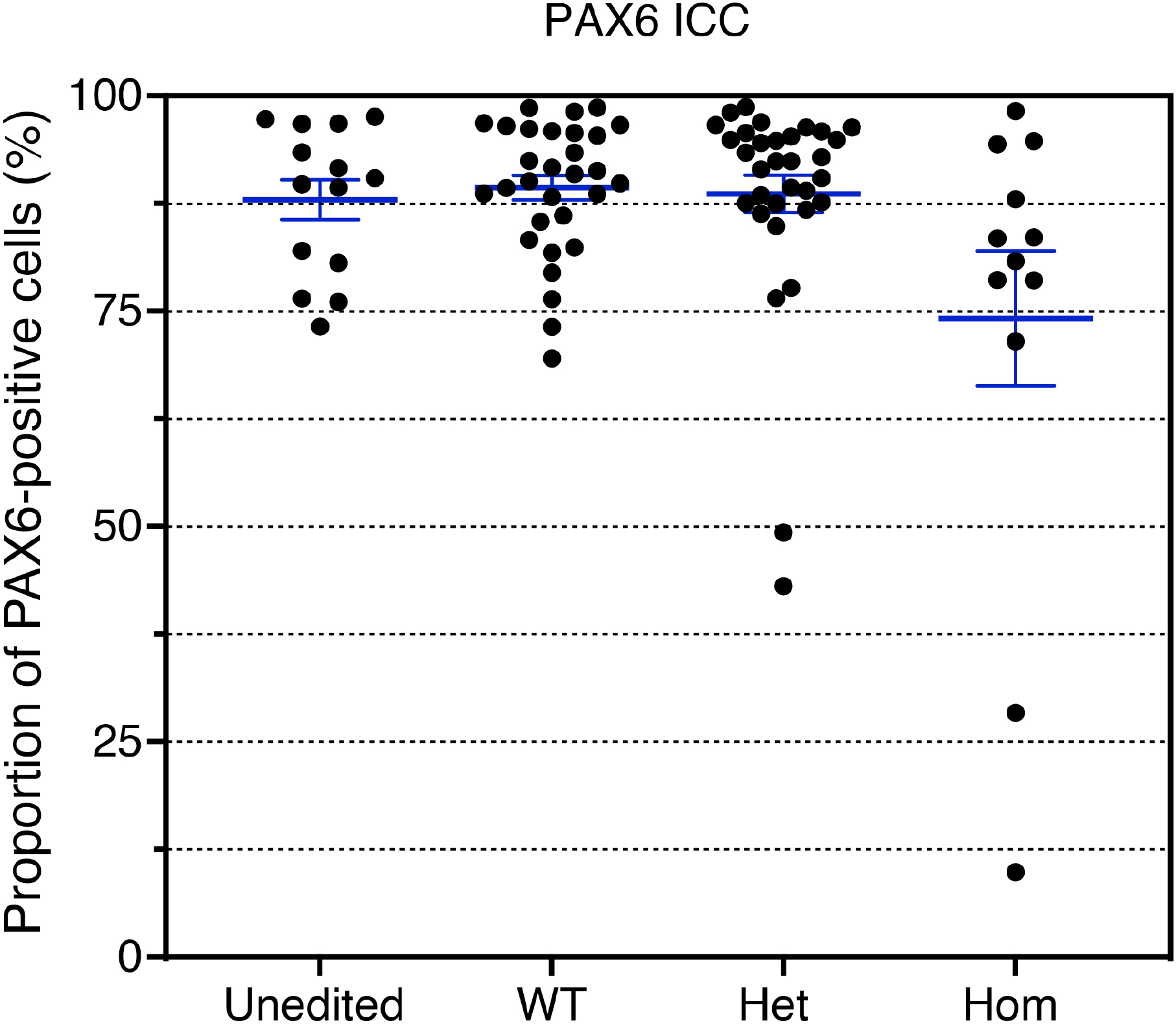
PAX6 expression rates across gene sets. The distribution of PAX6 expression rates across 93 cell lines from 16 gene sets successfully differentiated into neural stem cells in 2018 and 2019. Data is separated into Unedited, wild-type (WT), heterozygous (Het) and homozygous (Hom) mutant lines, and shows that the vast majority of cell lines display greater than 80% PAX6-positive cells. Bars denote mean +/- SEM.

**Figure 5.**
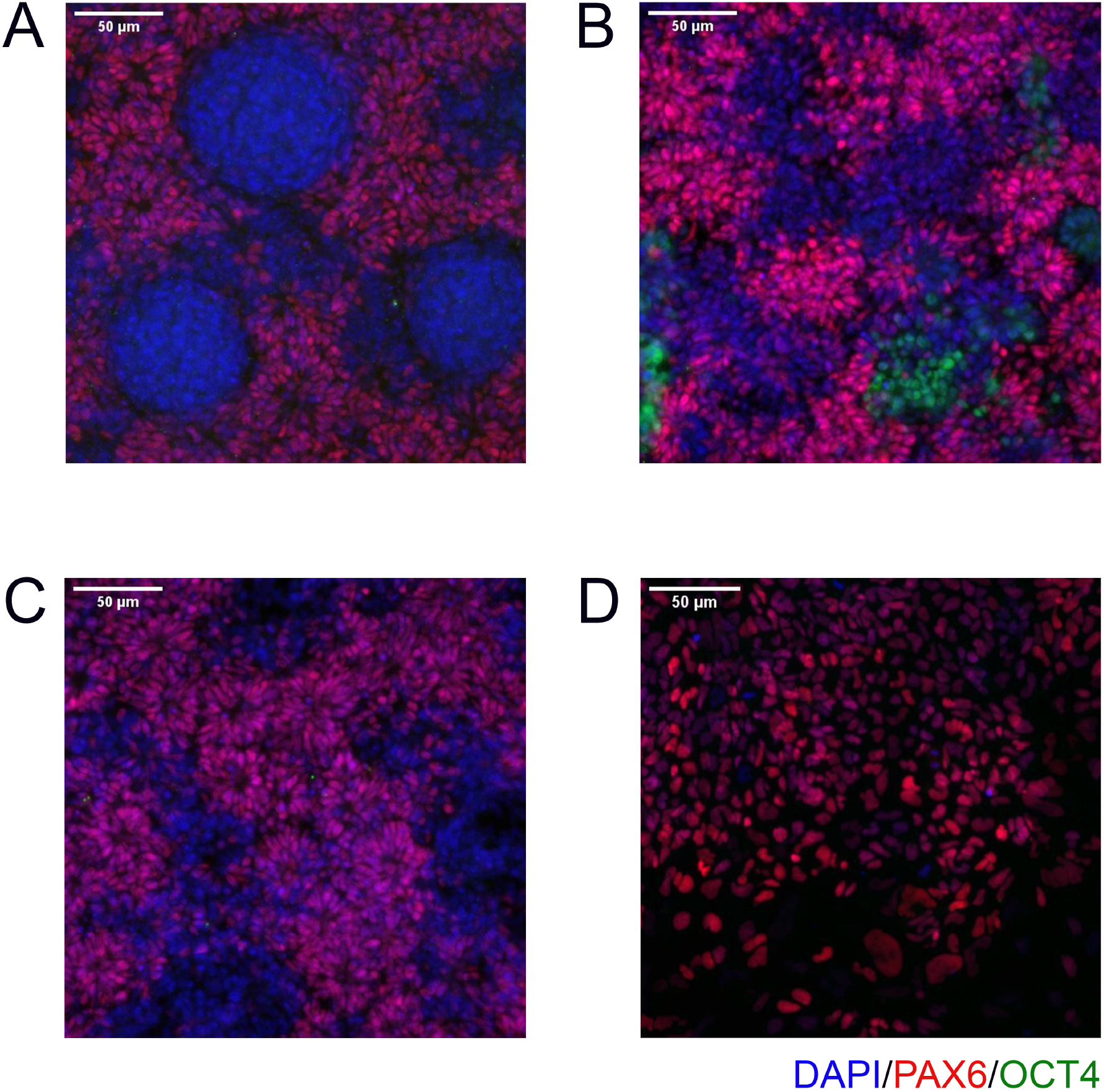
Examples of common issues seen in immunocytochemistry. **(A)** circular clumps of cells that do not express PAX6, surrounded by PAX6-positive cells. **(B)** Patchy PAX6 expression, with areas also expressing OCT4. **(C)** Similar to **(B)**, Patchy PAX6 expression, however in this case the morphology is still good with obvious neural rosettes. **(D)** NSCs have failed to properly attach to the 96 well plate and so do not take on the expected morphology. Scale bar, 50μm (20x).

### Troubleshooting Immunocytochemistry

#### Potential Problem 1

Poor attachment of NSCs to the 96-well plate (Figure 5D) or detachment after fixing or staining. Poor attachment can be caused by incorrect seeding density; ensure that the correct volume of NSCs is isolated for ICC during the Day 10 passage, and that the cells are thoroughly re-suspended before plating. Detachment after fixing or staining can usually be avoided by dispensing liquid slowly onto the sides and making sure to avoid touching the bottom of the wells when aspirating. In the case of a high degree of cell detachment, additional ICC data can be obtained without repeating the differentiation, by thawing the cryopreserved NSCs directly onto the 96-well plate, culturing until confluent (4-5 days), then fixing and staining as before.

#### Potential Problem 2

iPSCs become overcrowded on the 96-well plate. iPSCs colonies should be present, as opposed to a monolayer. Overcrowding is likely due to seeding too many cells on Day 10, either through performing the split ratio incorrectly, seeding the incorrect volume, or not re-suspending the cells thoroughly before seeding. It may also be due to culturing the cells for too long before fixing (3-4 days is usually optimal).

### Neural Stem Cell Thawing and Passaging Results

After completing the neural differentiation, a vial of each cryopreserved NSC sample is thawed in order to confirm that the frozen cells are viable. One vial is thawed to one well of a 6-well plate, which is expected to form a confluent monolayer by the following day. 3-4 days later, the cells should appear as a densely packed layer of cells, and are ready to passage (Figure 6, top two panels). They may also display neural rosettes at this stage, as shown in Figure 6. After passaging 1:1, they should form a confluent monolayer again within 1-2 days, with little change in the culture compared to before passaging (Figure 6, bottom two panels).

**Figure 6.**
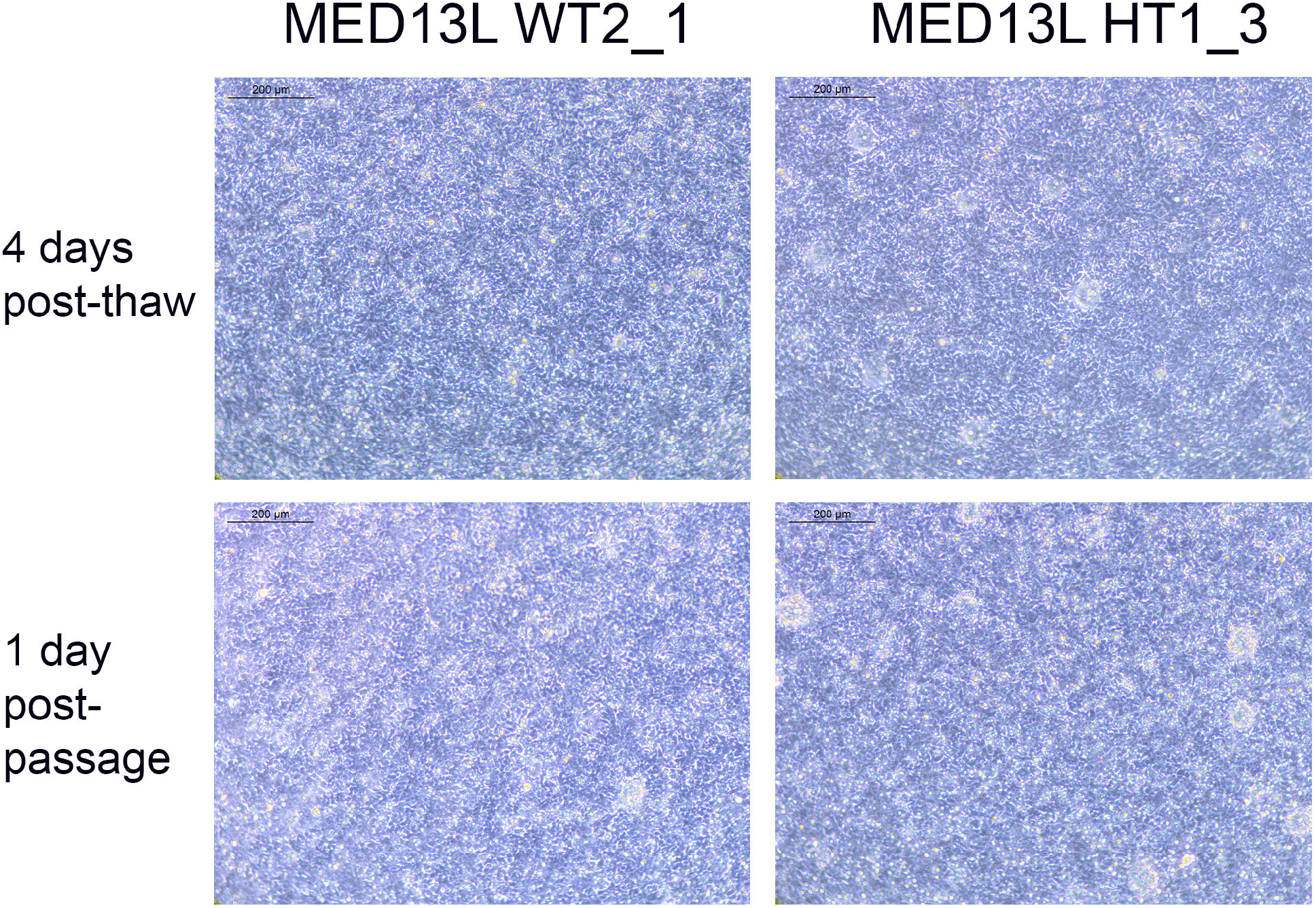
Neural Stem Cell QC images. Brightfield images of two cell lines from the MED13L gene set (WT2_1, and HT1_3), which both display typical morphology. The top images are from 4 days after the QC thaw (just before passage), and the bottom two images are from the following day, 1 day after a 1:1 passage. Scale bar, 200μm (10x).

### Troubleshooting Neural Stem Cell Thawing

#### Potential Problem 1

NSCs have not formed a confluent monolayer, or have a flat, uncompacted morphology 4-5 days after thawing (Figure 3F). This is likely due to an error during the cryopreservation process, which led to too few cells being frozen per vial. It may alternatively be due to poor viability after thawing, which can occur if the vial is left in the 37 ºC water bath for too long, or as a result of being left in the DMSO-containing Neural Freezing Medium for a prolonged period after the cells have thawed.

#### Potential Problem 2

NSCs do not form a confluent monolayer after passaging, or appear clumpy. The absence of a confluent monolayer will usually indicate that not enough cells were seeded back down after passage. This can be due to insufficient accutase treatment during the passage, or insufficient resuspension after centrifugation. It may also help to filter the cells with a 70µM filter after accutase treatment, to remove larger clumps of the cells.

### Additional Validation Data

The following results are included to demonstrate the robustness and specificity of our neural differentiation protocol. They include data from experimental processes that are not routinely part of the regular pipeline.

### ICC compared to flow cytometry

To validate the ICC results, a side-by-side comparison with flow cytometry for the same cell lines was performed (Figure 7). The results for two cell lines from the SETD1B gene set (WT1 and HT1), and the parental Kolf2C1_WT NSCs and iPSCs are shown. ICC images are displayed in Figure 7A and 7B, while the graphs in 7C and 7D show a direct comparison of the quantitative expression data gathered by the Cellomics Arrayscan and flow cytometry results. The NSCs used for both experimental methods were thawed simultaneously, pooled and seeded to a 96-well plate for ICC, and a 6-well plate for flow cytometry. Both were cultured for 4 days, before fixing and staining the cells for ICC, and pelleting, fixing and staining the cells for flow cytometry. The same primary and secondary antibodies were used for both processes. These data (Figure 7C, D) demonstrate the high concordance between both experimental methods, reflected by the similarity in the measured expression of PAX6 and FOXG1. The measured expression levels vary dramatically between the cell lines, with the heterozygous line exhibiting much reduced expression of both markers. This is also clear from the ICC images (Figure 7A and B), in which the morphology of the cells is abnormal and that there is a visible lack of PAX6 and FOXG1 staining in many of the cells. As expected, the iPSCs display no visible expression of either NSC marker, but do express high levels of OCT4. The flow cytometry data confirms the lack of NSC marker in the iPSCs. Flow cytometry plots can be found in Supplementary Figure 1.

**Figure 7.**
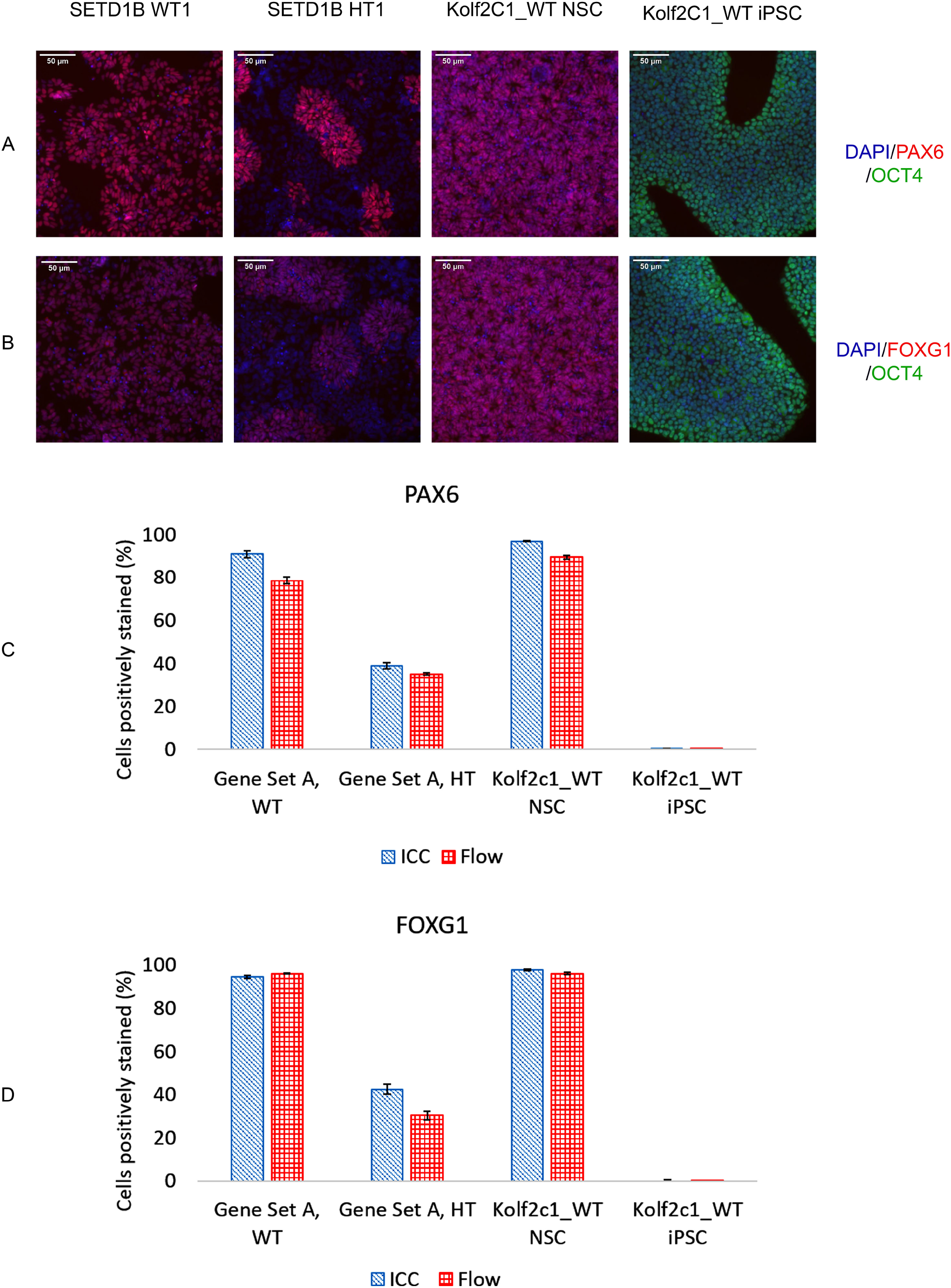
immunocytochemistry (ICC) agrees with flow cytometry results. Two cell lines from the SETD1B gene set (SETD1B WT1 and SETD1B HT1), the parental Kolf2C1_WT NSCs and iPSCs were thawed and simultaneously seeded on a 6-well plate for flow cytometry, and a 96-well plate for ICC. (**A**,**B**) Representative composite ICC images for PAX6/OCT4 expression (**A**) and FOXG1/OCT4 expression (**B**). (**C**,**D**) Quantification of samples using the Cellomics Arrayscan (ICC, blue bars) and the flow cytometer (red bars) for PAX6 expression (**C**) and FOXG1 expression (**D**). Scale bars, 50μm (20x).

### RT-qPCR using MED13L gene set

RT-qPCR was performed on differentiated NSCs from the MED13L gene set, containing two wild-type and two heterozygous lines, in order to measure the fold-change in expression in the NSCs compared to undifferentiated Kolf2C1_WT iPSCs. Each differentiation is performed as two technical replicates, but for simplicity, only one of each was used for RT-qPCR. The differentiation was performed twice using the same cell lines, once in 2019 and once in 2020. Supplementary Figure 2 shows the results for these experiments, with 2019 data displayed in blue and 2020 data in orange. Markers of neural stem cell identity – *SOX1, PAX6, NESTIN, FOXG1* and *GFAP* – all increase in every NSC cell line. As expected, the increase in NSC marker expression is accompanied by a loss of *POU5F1 (OCT4)* expression, confirming that the cells lose their pluripotency as they differentiate. Broadly, these data show that neural stem/precursor cell identity is achieved through our protocol, and in this particular experiment, is not prevented by heterozygous knockout of MED13L.

### RNA-Seq

We performed RNAseq on the iPSCs and differentiated NSCs of wild-type lines of the MED13L gene set (Figure 9). We observed a strong upregulation of neural stem cell markers (Figure 9A, *PAX6, SOX1*) with the expression of region-specific markers (Figure 9B, forebrain and cortex-specific markers: *FOXG1, OTX2, LHX2, SIX3*, and *EOMES/TBR2*), and minimal expression of mid and hindbrain markers (*GBX2, IRX3*). As expected, there was a concomitant loss of iPSC markers (Figure 9C, *NANOG, OCT4/POU5F1*), demonstrating a successful differentiation. Figure 10 shows the correlation of expressed genes between technical replicates for MED13L, demonstrating the high consistency between independently differentiated replicates of samples from the same clonal source. These data support our ICC and RT-qPCR results, and confirm successful regionalisation of our NSCs during differentiation.

**Figure 8.**
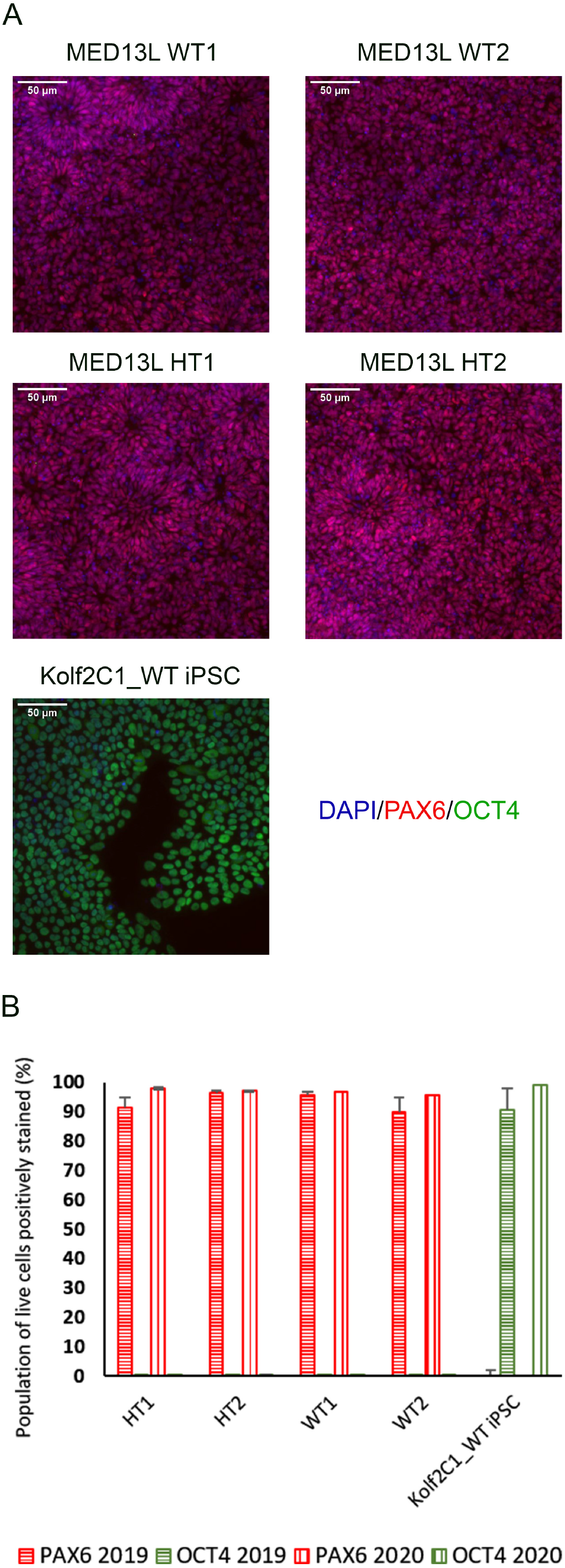
Gene expression analysis for the MED13L knockout gene set. **(A)** Immunocytochemistry images of differentiated NSCs from the MED13L gene set, plus Kolf2C1_WT iPSC control sample, showing expression of PAX6 and OCT4. **(B)** Quantification of immunocytochemistry results, comparing data from two independent rounds of differentiation (2019 and 2020).

**Figure 9:**
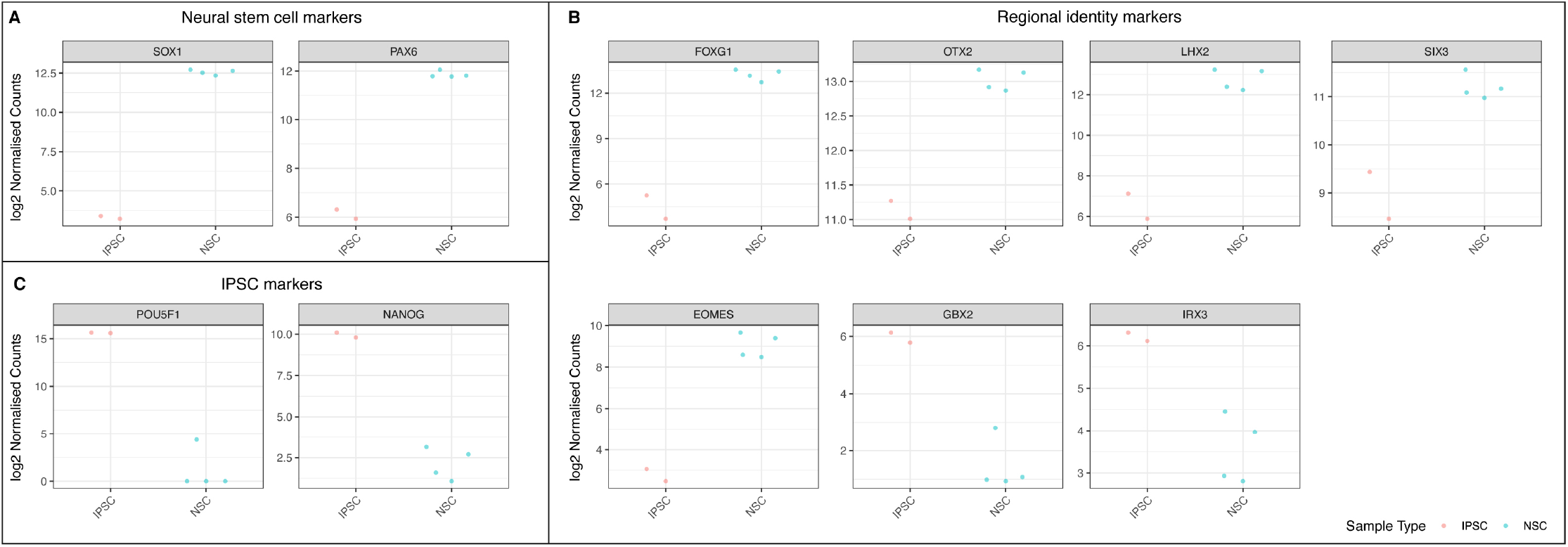
RNA-Seq analysis of the wild-type cell lines from the MED13L gene set, comparing undifferentiated iPSC lines to differentiated neural stem cells. Log2 normalised read counts of **(A)** two neural stem cell markers, **(B)** 7 markers of regional NSC identity **(C)** two pluripotent stem cell markers. Note that there are four samples for neural precursor cells and two iPSC samples, because each cell line was split into two technical replicates immediately prior to differentiation.

**Figure 10:**
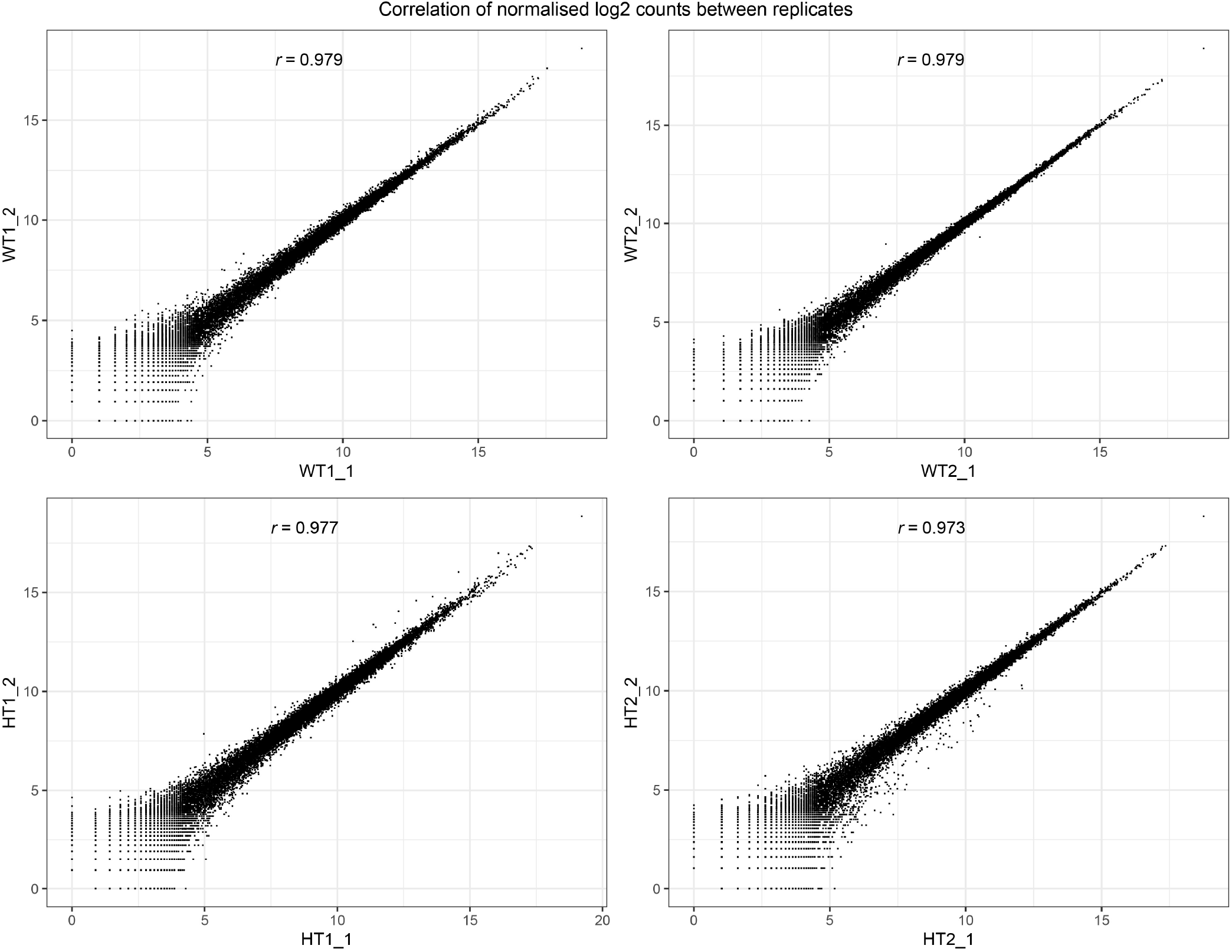
Consistency of RNA-Seq replicates. Correlation between levels of gene expression (normalised Log2 counts) of differentiation replicates of neural precursors for the MED13L gene set, with Pearson correlation scores for each comparison.

### Differentiation of iPSCs from different donors

To demonstrate that our differentiation protocol can generate NSCs from a variety of iPSC lines, it was tested using three lines from the HipSci resource (https://www.hipsci.org) (Pelm_3, Podx_1 and Sojd_3) in parallel with our standard Kolf2c1_WT. The standard neural differentiation protocol was followed, as described in section 3.2 – 3.6. All lines successfully completed the differentiation, and were subsequently stained using the protocol described in section 3.9. Cell pellets were collected at Day −1 and Day 14 to carry out RT-qPCR. Images of the iPSC lines before starting neural induction (Day −1) are shown in Figure 11A. There is some variation in the resulting morphology of each line at Day 14; however, all are acceptable NSC morphology. Figure 11B shows the ICC images for each line, and Figure 11C shows the quantification of PAX6-positive and OCT4-positive cells. All NSC lines are >90% PAX6-positive and do not express OCT4. Together, these experiments demonstrate that our neural differentiation protocol can successfully produce neural stem cells from iPSCs derived from different donors. This was confirmed by RT-qPCR (Supplementary Figure 3), which shows an upregulation of all NSC markers tested, in each NSC line (relative to the equivalent, undifferentiated iPSC line). Kolf2C1_WT has the largest increase in *PAX6, FOXG1* and *NESTIN* expression, which may reflect the fact that the differentiation protocol was optimised using this cell line: Further adjustments may be required to achieve the optimal yield and purity of NSCs in other genetic backgrounds. We would recommend adjusting the length of the neural induction period and ensuring high quality of the iPSC population at induction, as a starting point for this optimisation, as these parameters have been shown to have an impact on the protocol outcome. Nonetheless, our standard 14 day protocol yielded high purity NSCs with no residual iPSC contamination. The robustness and transferability of our protocol was further confirmed by the differentiation results of two additional lines (kucg2 and paim1), completed by an independent group at a different facility (work completed by the Kilpinen research group at UCL (Supplementary Figure 4)), successfully producing PAX6-positive lines in a 14 day protocol run.

**Figure 11.**
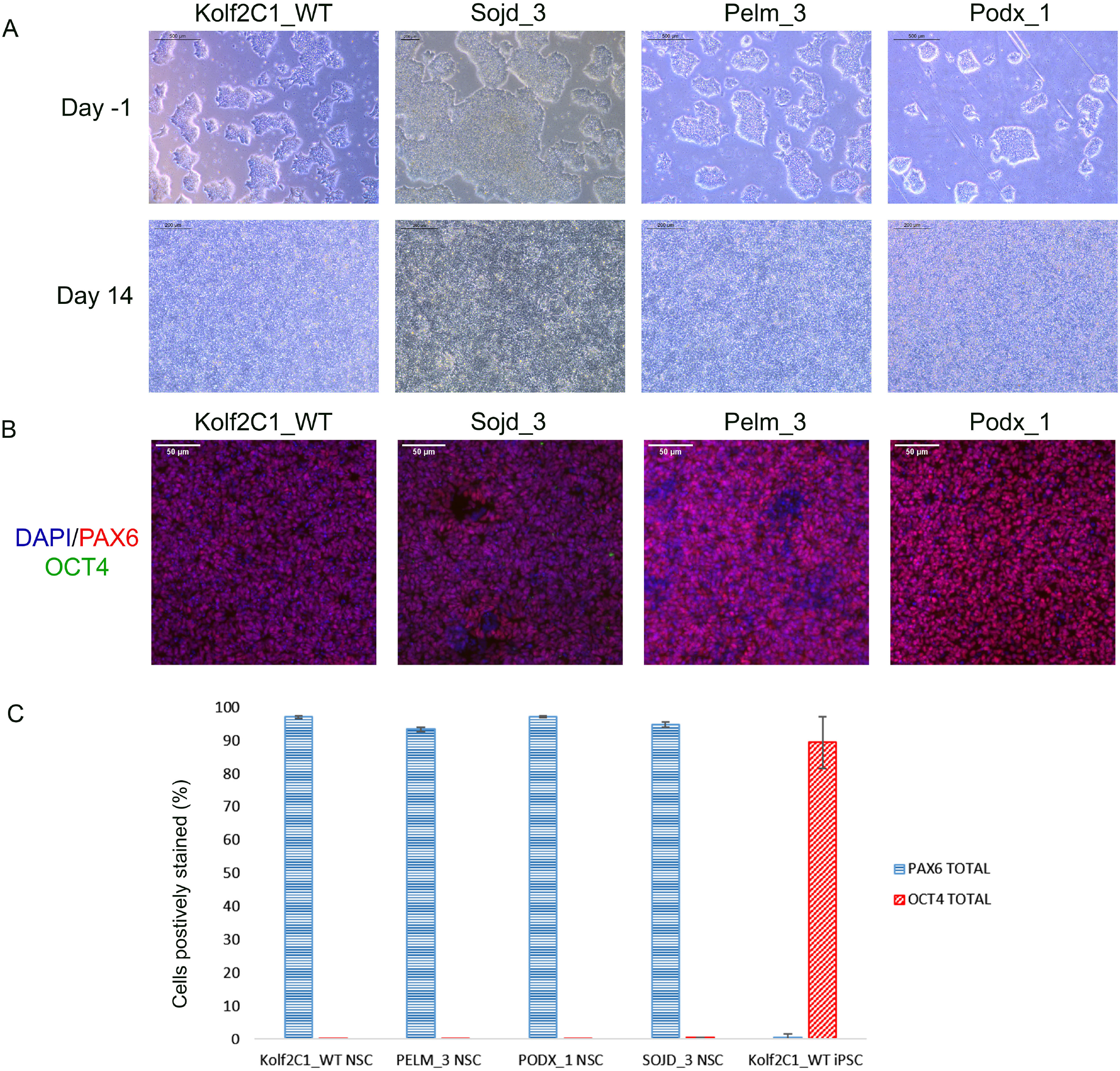
Differentiation outcomes of additional iPSC lines. **(A)** Brightfield images for Kolf2C1_WT, Sojd_3, Pelm_3 and Podx_1 on Day −1 prior to the Neural Induction passage (Scale bar 500μm, 5x), and on Day 14 prior to cryopreservation (Scale bar, 200μm, 10x). Note that Sojd_3 Day −1 image has a different sized scale bar (200μm) but the magnification is the same. **(B)** Immunocytochemistry images showing PAX6, OCT4 and DAPI staining of NSCs generated from all four iPSC lines (Scale bar 50μm, 20x). **(C)** Quantification of immunocytochemistry results, displaying the percentage of cells positively stained with PAX6 and OCT4 for each cell line.

## Discussion

In this paper, we have described a robust method to generate cortical neural stem cells from both unmodified and CRISPR-edited hiPSC lines. Using this method, we have successfully differentiated over 100 cell lines, validated by RNA sequencing and ICC assays of NSC marker expression levels, demonstrating that this method can be used for high-throughput gene expression studies. We use a dual-SMAD inhibition approach, an established method for generating neural precursors from pluripotent stem cells, but additionally include the Wnt pathway inhibitor XAV939 in the neural induction medium, as it has been shown to increase the number of forebrain-specific FOXG1-positive neural precursors (Maroof et al, 2013). Development of a robust, reproducible method for generating NSCs is essential in establishing a reliable model for studying development *in vitro*.

The protocol described in this paper has been subject to various refinements and improvements over a number of years. The most notable of these was the shortening from a 20-day protocol (including an extra 6 days of NSC expansion) to the 14-day protocol described here. The extended expansion phase, while achieving the intended increase in cell numbers, also increased the risk of bacterial contamination and spontaneous differentiation. In addition, a variety of plate coatings were tested, including full length vitronectin, laminin and laminin combined with poly-L-ornithine (data not shown), and it was found that these substrates did not provide the right adhesive environment for a homogeneous cell population, whereas recombinant vitronectin (VTN-N), diluted to 1:50, permitted the formation of a uniform NSC culture.

Our efficient, scalable protocol is currently being implemented in a large-scale project to examine the transcriptional consequences of loss-of-function in dozens of developmental disorder associated genes, relying on CRISPR-engineered isogenic disease model lines (for example Levitin et al., 2022). Our current protocol allows a single researcher to easily handle the simultaneous differentiation of as many as 12 individual cell lines, generating 2-4×10^7^ NSCs per line at the end of the 14-day protocol. This allows for numerous downstream assays to be performed, such as RNAseq, flow cytometry, and RT-qPCR. Importantly, we find a high consistency in our protocol, as demonstrated by the successful differentiation of dozens of cell lines over a 2-year period, during which the vast majority of generated cell lines showed more than 80% PAX6-positive cells. RNAseq analysis reveals a strong correlation between technical replicates, indicating robustness of the protocol against experimental variation. Variation between individual differentiation runs and between cell lines is commonly seen and may be attributable to the knockout mutations, or slight variations in culture conditions (for example, media temperature and time since preparation). In addition, different cell lines may vary in the time taken to reach the NSC stage. In the context of our large-scale differentiation pipeline, we prioritise the consistency of the method through use of a standardised procedure, which could ultimately lead to less consistent outcomes. Depending on the context of the project and experimental design, it may be necessary to have more flexibility in the methods used, and adjust the duration of neural induction according to the observed morphology (or additional analytical methods). Similarly, variation in morphology was seen when a team at a different organisation tested the protocol, but importantly all cell lines show the expected levels of PAX6-positive cells.

Our protocol uses an adherent cell culture system to generate NSCs from human iPSCs on a large scale, but other approaches may be capable of achieving similar results. Zhang et al (2001) found that when neural differentiation occurs from an embryoid body (EB) starting point, the differentiation process tends to recapitulate early *in vivo* development through the formation of neural tube-like structures, which contain neural rosettes. The EB itself is similar to the early embryo, with mature EBs mirroring embryonic blastocyst structure (Liyang et al, 2014). It has been argued that the EB model has greater apparent similarity to the *in vivo* process and is therefore more physiologically relevant. However, EB culture can lead to the presence of a diversity of microenvironments that are less well controlled, potentially generating a more heterogeneous cell population or more variable outcomes. Furthermore, the size of the EB is a crucial determining factor in the outcome, thus introducing additional opportunities for undesirable end-point variation (Bauwens et al, 2008). Other approaches to neural differentiation that avoid an EB culture step include those using floating aggregates known as neurospheres (Zhou et al, 2016). However, these are vulnerable to some of the same issues as EB culture, as the cell density of each neurosphere will alter the microenvironment, leading to differential positional cues and proliferation capacity (Jensen & Parmar, 2006). Moreover, this system is usually not amenable to large-scale studies, due to difficulties in maintaining consistency in neurosphere size and quantity across a large number of experimental samples (D’Aiuto et al, 2015).

The use of monolayer culture, as we have demonstrated, offers a highly controlled microenvironment, ensuring uniform cytokine exposure across the cell population. Ensuring the correct cell density at neural induction is important, as low plating density combined with dual-SMAD inhibition encourages the formation of PAX6-negative neural crest-like cells, whereas high density results in a near-exclusive PAX6-positive cell population (Chambers et al, 2009). In the vast majority of cases, the entire cell population at the end of our 14-day protocol is PAX6 positive, and therefore can all be considered neural precursors. Another key advantage to monolayer culture is the ability to visually monitor the cells as they differentiate. This is in contrast to EB/neurosphere approaches, in which NSCs may emerge within the centre, but cannot be readily identified in brightfield, but only after fixing and staining for cell-type specific markers or re-plating into a 2D culture system. Our protocol allows for continuous observation, generates consistent results and importantly, takes only 10 days to reach a high purity neural stem cell state, followed by a 4-day maintenance period in which neural rosettes emerge.

The neural differentiation protocol described by Shi et al (2012) is similar to our approach in several ways, including the use of a confluent iPSC monolayer as the starting point for neural induction, and maintaining 100% confluence throughout differentiation. However, our protocol relies on single-cell accutase dissociation, allowing us to more accurately count and seed the cells at the same density every time. This ensures consistency within and between experiments. Our protocol ends with cryopreservation of the NSCs on Day 14, whereas Shi et al (2012) further expand the NSCs for an additional 10-15 days, adding bFGF to promote their expansion, and then optionally cryopreservation or continuing to culture the cells until mature cortical neurons form. They do not suggest cryopreservation until neurons begin to appear around the rosettes, approximately 20-30 days into the protocol. In contrast, we prefer to collect and freeze cells at an early time point, when cellular heterogeneity is more limited. Our current project analyses RNA-seq at Day 14, a time point at which we can robustly identify differentially-expressed genes as a result of knockout mutations. Longer culture and further differentiation would allow a researcher to examine more mature and differentiated neuronal and glial derivatives. We always perform a QC thaw of the cryopreserved NSCs, in order to confirm their viability and demonstrate that they can be cultured for a longer period after thawing, if desired. Preliminary experiments (data not shown) confirm that further culturing and differentiation of our cryopreserved Day 14 NSCs can generate a diversity of cortical neurons, thus providing a route to varied downstream analyses of our disease models.

While our protocol is amenable to differentiating CRISPR-edited iPSC lines, there is the potential for the differentiation process itself to be impacted by the mutations introduced, given that many of the genes we study are involved in neurodevelopmental processes. There have been occasions where we concluded that the engineered mutation has prevented us from achieving a homogenous NSC population, although the robustness of the neural induction is such that this is a rare event. Distinguishing a technical failure of the protocol from one caused by genotype differences can be challenging and we rely on consistency between biological (i.e. different cell lines of the same zygosity or genotype) and technical (i.e. independent wells with the same line) replicates in the first instance. More detailed analysis of the resulting RNA-seq may help to identify consistent genotype-related expression changes, and potentially clarify the nature of the problem, or the alternate cell type generated.

The use of ICC as a quality control method provides us with a quick and convenient way of quantifying NSCs in our cultures. While we currently use three ICC markers (PAX6 and FOXG1, and OCT4), one could modify the scale of cell plating to accommodate additional markers. The 96-well plate format and the use of the Cellomics Arrayscan allow us to image the cells efficiently, gaining both qualitative and quantitative data for many cell lines at once, providing a less intensive workflow than using a standard fluorescent microscope. In addition, the Cellomics is able to give highly concordant quantification of expression when compared to flow cytometry. Overall, the Cellomics is very effective in our workflow for deciding whether a differentiation has been successful, and it is a precursor to the RNA-Seq analysis that provides a much more in-depth, sensitive view of gene expression.

To understand the molecular consequences of our gene knockouts on NSC differentiation, we routinely use bulk RNA-seq followed by differential gene expression analysis. This provides an unbiased, sensitive method for identifying changes in gene expression in our disease models, insights into alterations in various biological pathways, and the establishment of “transcriptional signatures” that can be used to compare between samples, diseases, and against the signatures of potential therapeutic agents. The relatively homogenous nature of the cell populations in our cultures make this bulk approach effective. Changes in cellular subtypes or population proportions may be missed by such an approach, which can be complemented by single cell RNA-Seq methods: These allow the identification of individual cell identities and proportions, at the expense of sensitivity to detect more lowly expressed genes. Applying a combination of approaches (e.g. ATAC-Seq, mass spectrometry) would permit a more exhaustive examination of the molecular underpinnings of neurodevelopmental disorders, with a concomitant increase in complexity and cost. In this study, we used RNA-Seq in its most basic form, to generate gene expression data highlighting the successful differentiation and regionalisation of our cortical NSCs.

Importantly, we show that our differentiation protocol will work for a number of different iPSC lines. We demonstrate that our protocol is reproducible by replicating these results in an independent lab at a different institute, highlighting that similar results can be achieved but not without a certain level of variation, between cell lines and between sites. We have provided a detailed version of our differentiation protocol, with the hope that other laboratories will find similar success in generating neural stem cells from iPSCs. We additionally provide the link to our Protocols.io webpage, which describes our ICC assay in further detail.

Our academia/industry partnership project aims to provide gene expression data for hundreds of knockout NSC lines, ultimately aiming to find similarities in the aberrant gene expression between different developmental disorder models. This may help to find common therapeutic targets, which can then be investigated further, offering hope to the many families living with rare developmental disorders worldwide. The scalable approach we describe is applicable in many other contexts, for example in high-throughput drug screening, which requires high levels of consistency and low batch-to-batch variation. The defined 10-day NI period helps to standardise the differentiation process, reducing complexity, and we have shown that our protocol will work for a large number of knockout and wild-type iPSC lines. Importantly, iPSCs can be expanded continuously, providing a limitless quantity of cells. These can be edited using techniques such as CRISPR/Cas9, or reprogrammed directly from samples taken from patients, in order to establish a disease model. They can be further characterised and studied in depth, and potentially identify whether the disorder affects NSC differentiation or maintenance, and can subsequently be used for drug screening or further differentiation. In the context of neural differentiation, there are broad applications, stretching from neurodevelopmental disorders to dementia.

## Abbreviations

NIM: Neural Induction Media
DPBS+/+: Dulbecco’s Phosphate Buffered Saline with MgCl_2_ and CaCl_2_
NMM: Neural Maintenance Media
iPSC: induced pluripotent stem cell
NSC: Neural Stem Cell
EB: embryoid body

## Acknowledgements

The neural differentiation protocol has previously been published online at: https://www.protocols.io/view/differentiation-of-human-induced-pluripotent-stem-bgbxjspn.

We would like to thank all the members of Cellular Generation & Phenotyping team for their support and guidance, and helping with weekend media changes. We would especially like to thank the previous members of the DDD-NeuGen Team (Emily Relton, Neophytos Kouphou, Rohinder Bains and Maria Imaz) for contributing to the development and refinement of the neural differentiation protocol. We would like to thank the Gene Editing team for creating all of the knockout iPSC lines, the Cytometry Core Facility for providing their equipment and expertise, Sample Management and DNA Pipelines for providing the RNA-Sequencing, and Vivek Iyer and Guillaume G. Noell for Bioinformatic support.

## Author Contributions

The manuscript was written and edited by AN with assistance from MHLA and SSG. The DDD-NeuGen study was conceived and designed by SSG and MEH. Knockout iPSCs were created by BN and TT. Pre-differentiation culture of iPSCs and differentiation into neural stem cells was carried out by AN, MHLA and LF, overseen by AD and AH. Neural differentiation protocols were originally designed by SSG, BN, TT, AH, MP and MGT, and subject to further refinement by AD, MHLA, AN and LF. RNA-Seq was performed by the Wellcome Sanger Institute DNA Pipelines team and analysed by OAA and SSG. The additional validation experiments were performed by AN, with the exception of the neural differentiation of kucg2 and paim1, which was performed by CR in the lab of HK.

## Data Availability Statement

RNA-Seq data is available from the European Nucleotide Archive (www.ebi.ac.uk/ena) as follows: wild-type IPSCs (ERS3517948, ERS3517949), wild-type NPCs (ERS3517935,ERS3517936,ERS3517937,ERS3517938), *MED13L* heterozygous NPCs (ERS3517939,ERS3517940,ERS3517941,ERS3517942).

