## Supplementary Figures for "Differentiation of human induced pluripotent stem cells into cortical neural stem cells"

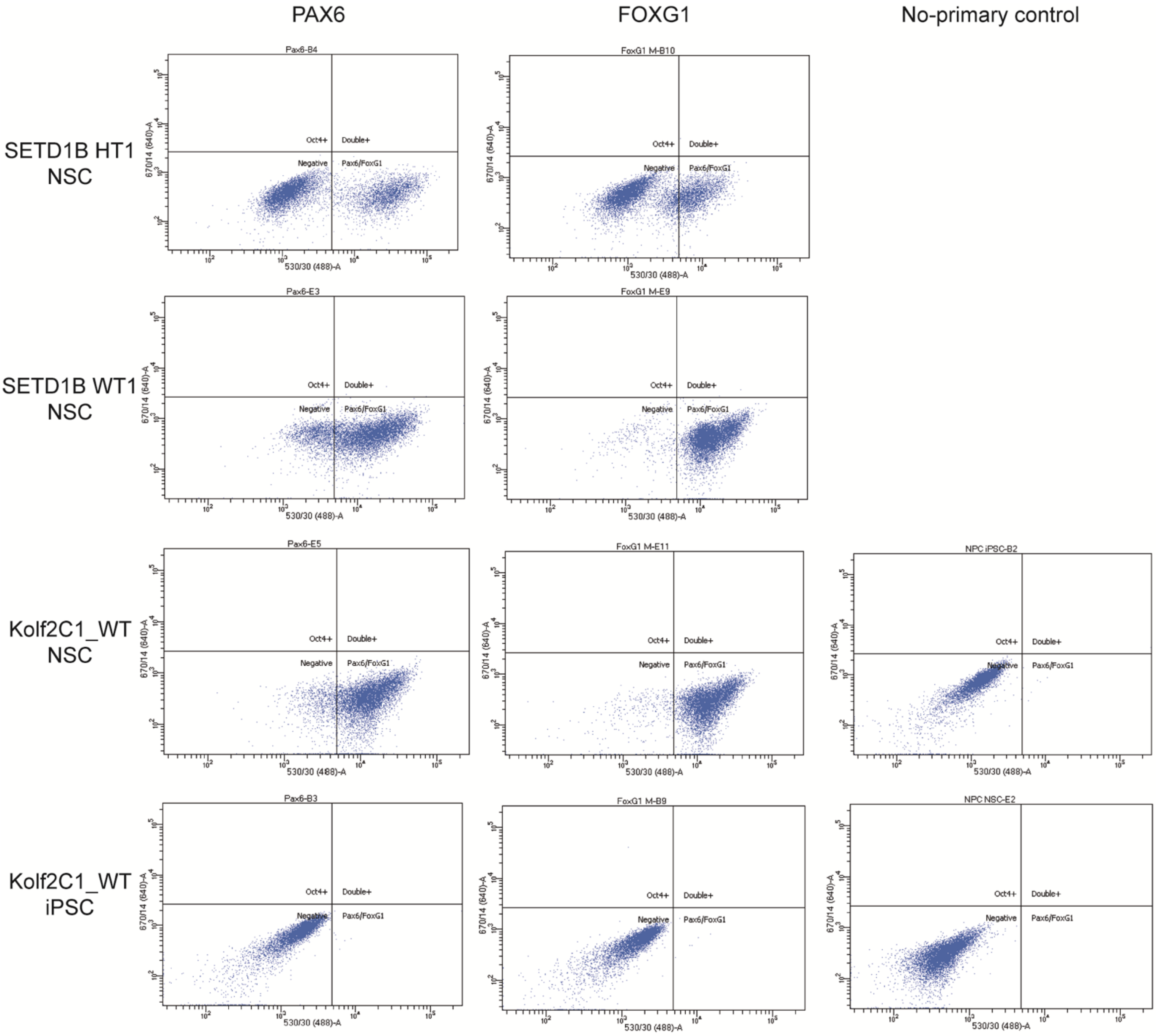


**Supplementary Figure 1. Flow Cytometry plots for NSCs and iPSCs.** Flow cytometry gating used to create the graphs displayed in Figure 7. iPSCs and NSCs stained using antibodies for OCT4 (y-axis) and either FOXG1 or PAX6 (x-axis), analysed on the BD Fortessa flow cytometer with BD FACSDiva. Adjustments were made to exclude doublets. Gating was set using the No-Primary controls for both Kolf2C1_WT iPSCs and NSCs and kept the same throughout all flow cytometry analysis. The first column shows the data for a heterozygous (HT) and a wild-type (WT) line from a gene set, plus Kolf2C1_WT NSCs and Kolf2C1_WT iPSCs, stained for OCT4 and PAX6. The second column shows the data for the same cell lines stained for OCT4 and FOXG1. The third column shows the data for the no-primary control samples (Kolf2C1_WT NSCs and iPSCs stained with only secondary antibodies).

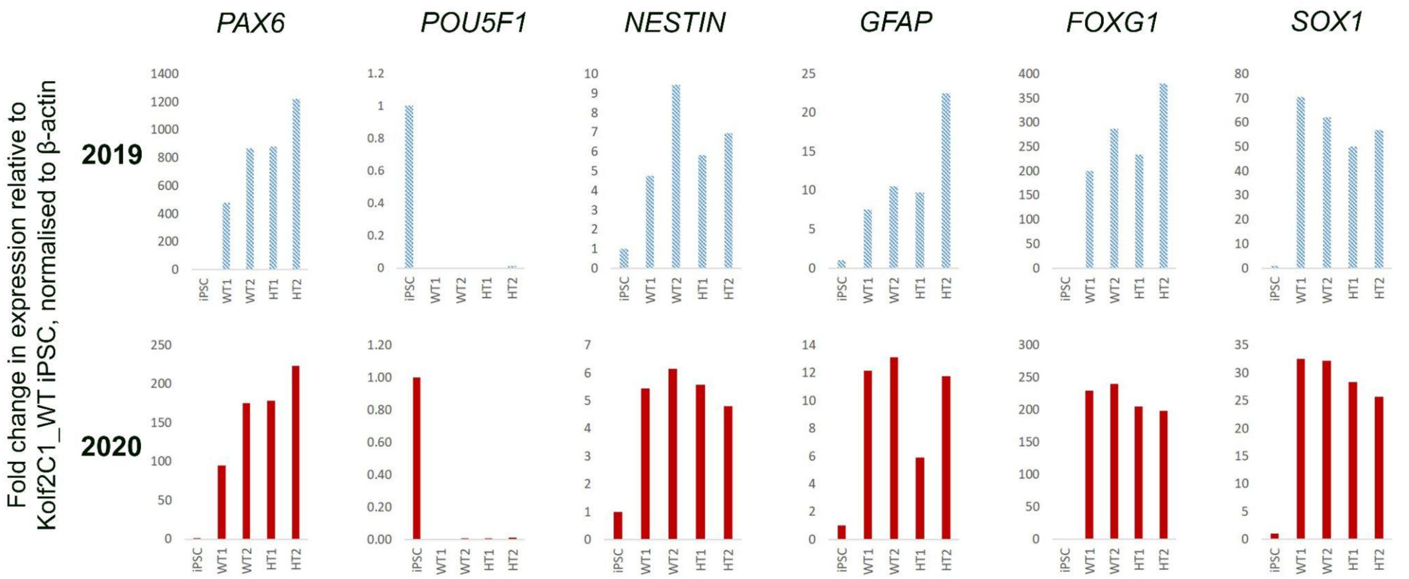


**Supplementary Figure 2. RT-qPCR gene expression analysis for the MED13L knockout gene set.** RT-qPCR data for the MED13L gene set, results derived from the same cell lines differentiated in 2019 (blue) and 2020 (orange). Analysis was performed with three technical replicates per sample. Results are displayed as fold change in expression relative to a Kolf2C1_WT iPSC sample, normalised to β-actin.

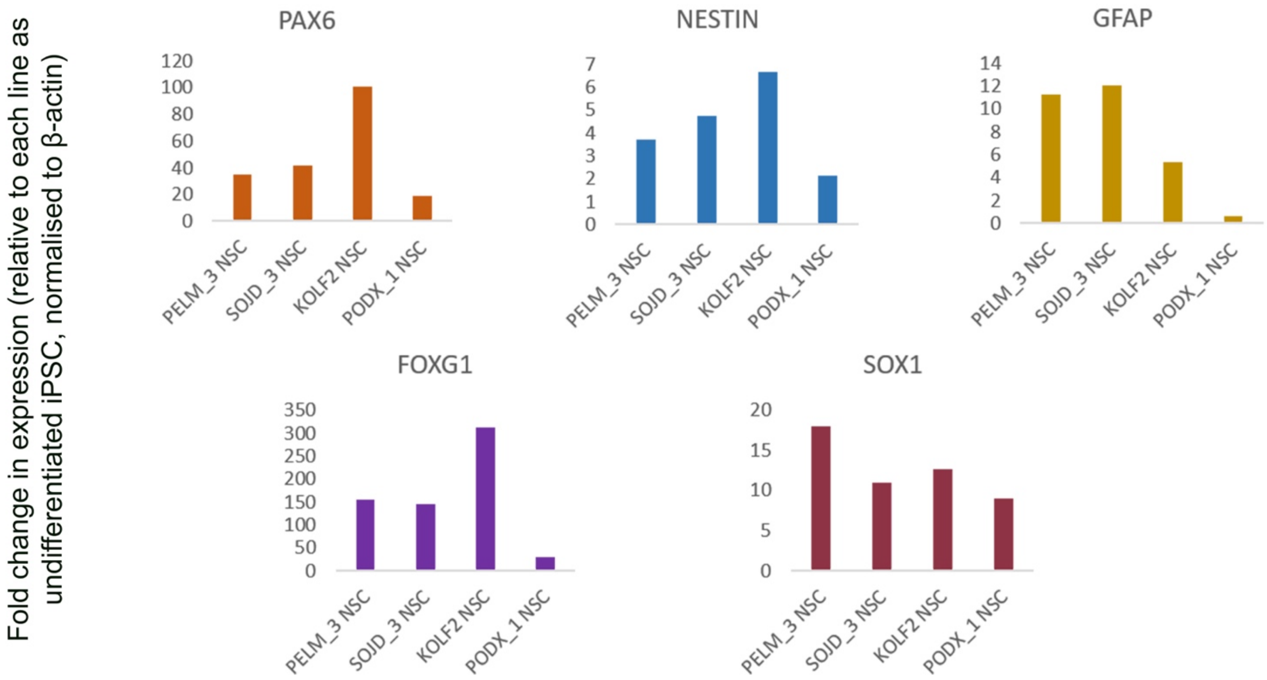


**Supplementary Figure 3. RT-qPCR gene expression analysis of additional differentiated iPSC lines.** RT-qPCR data showing the fold change in expression for each NSC line, relative to each cell line in its undifferentiated iPSC state, normalised to β-actin.

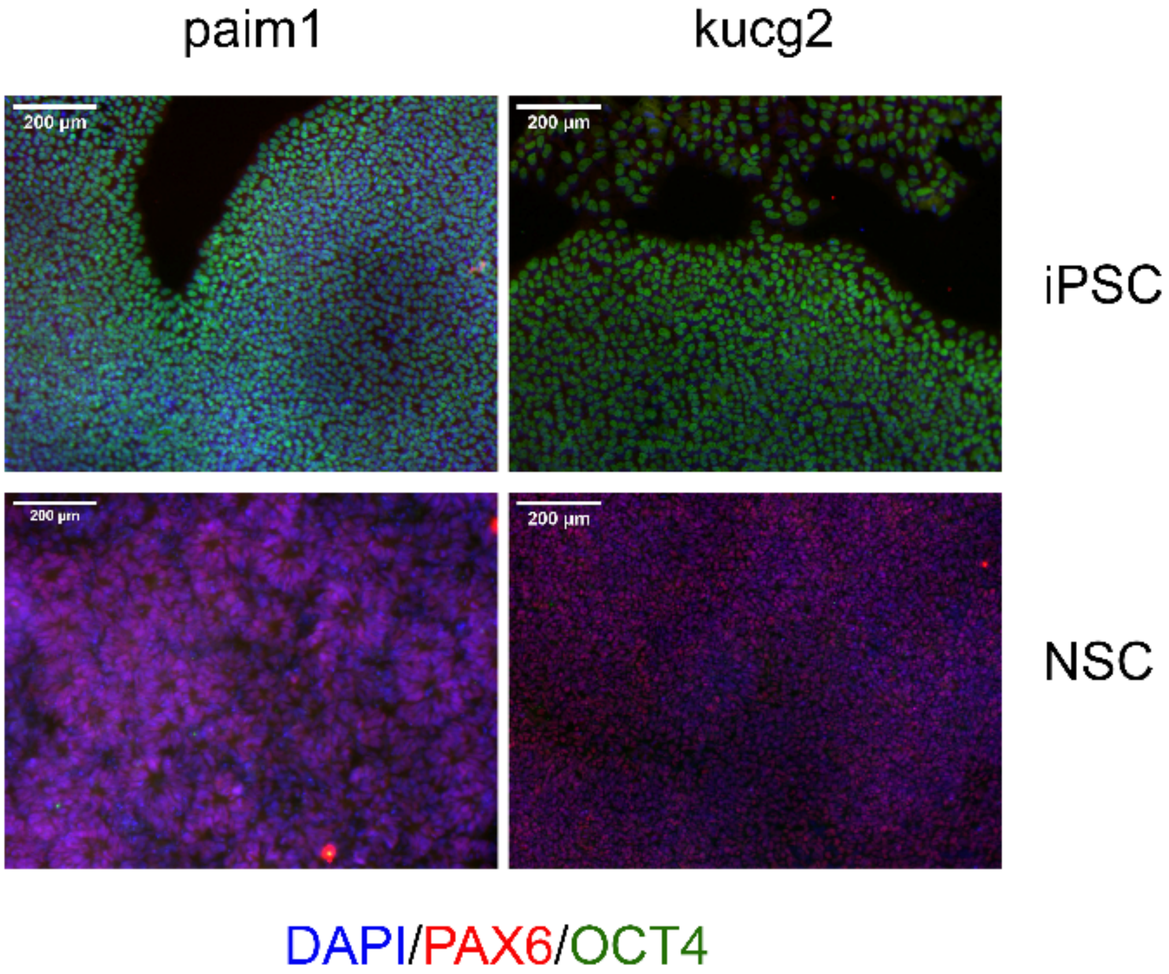


**Supplementary Figure 4. Successful differentiation of two additional iPSC lines.** Two iPSC lines (kucg2 and paim1) were differentiated into NSCs, then fixed and stained. The top panel shows the cell lines as iPSCs, and the bottom panel shows the same cell lines after NSC differentiation. Both cell types were stained for PAX6 and OCT4, plus DAPI. (Scale bar 200µm, 10x).
